# Cytoprotective and Neuroinductive Effects of Thiol-Containing Simple Signaling Molecules

**DOI:** 10.1101/2024.06.07.597935

**Authors:** Kylie J. Dahlgren, August J. Hemmerla, Marissa A. Moore, Daniela Calle, Bret D. Ulery

**Affiliations:** Department of Chemical and Biomedical Engineering, University of Missouri, Columbia, MO 65211, USA; Department of Biochemistry, University of Missouri, Columbia, MO 65211, USA

**Keywords:** cytoprotection, neuroinduction, simple signaling molecules, hydrogen sulfide (H_2_S), n-acetyl cysteine (NAC), glutathione (GSH), peripheral nerve injuries

## Abstract

Around 20 million Americans, mostly young adults, have suffered from a peripheral nerve injury due to trauma or a medical condition. However, prevailing treatments, including those that employ exogenous growth factors, often exhibit drawbacks outweighing their benefits, highlighting the need for novel solutions. Non-complex agents that govern bioactivity, termed simple signaling molecules (SSMs), offer an exciting alternative when compared to traditional approaches due to their small size, ready availability, and influence on cellular pathways. This study explores the cytoprotective and neuroinductive effects of three thiol-containing SSMs: hydrogen sulfide (H_2_S), n-acetyl cysteine (NAC), and glutathione (GSH). Neural stem cells (NE-4Cs) were exposed to toxic levels of hydrogen peroxide (H_2_O_2_) while supplemented with different concentrations of H_2_S, NAC, or GSH to assess their cytoprotective effects. To study the neuroinductivity of H_2_S, NAC, and GSH with NE-4Cs, immunofluorescence microscopy for an early neuronal marker, β3-tubulin, was employed to determine stem cell neural differentiation. Significant cytoprotection was found for NAC (2 - 8 mM) and GSH (2 - 12 mM), while H_2_S showed limited protective effects. Interestingly, the opposite result was observed for neuroinductivity as only H_2_S (7.75 - 125 μM) achieved a desirable effect while NAC and GSH had minimal impact on neural differentiation. This research establishes therapeutic concentration ranges of H_2_S, NAC, and GSH for future drug delivery systems targeting neural regeneration, especially for peripheral nerve injuries.

**LAY SUMMARY:** Approximately 20 million Americans, mainly young adults, suffer from peripheral nerve injuries, necessitating new effective treatments. Currently emerging approaches, including growth factor-based therapies, unfortunately face considerable limitations. This study explores the potential of simple signaling molecules (SSMs) - small, readily available agents - specifically hydrogen sulfide (H_2_S), n-acetyl cysteine (NAC), and glutathione (GSH) for their potential to be used to address issues underlying peripheral nerve injuries. Our findings reveal NAC and GSH have cytoprotective effects, whereas H_2_S is neuroinductive. This research identifies optimal concentrations for future drug delivery targeting neural regeneration, especially beneficial for peripheral nerve injuries.

**FUTURE WORKS:** Upcoming investigation will concentrate on exploiting the potential of novel SSM-releasing monomers within degradable polymeric systems. This involves defining therapeutic windows and optimizing polymer chemistry to induce desired cytoprotective and neuroinductive effects in neural stem cells, crucial for peripheral nerve injury repair. Emphasizing controlled release, this approach holds promise for developing advanced therapeutic strategies utilizing H_2_S, NAC, and/or GSH in nerve guidance conduits.

**GRAPHICAL ABSTRACT:** 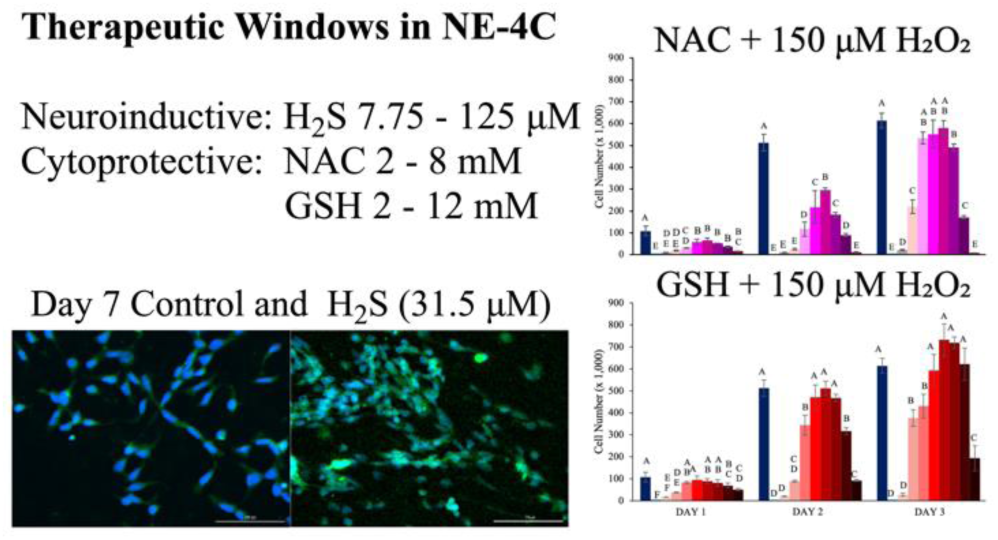

## INTRODUCTION

Damage to the nervous system is a considerable issue that significantly impacts patient well-being and outlook. Regardless, if damage occurs in the central or peripheral nervous system, it carries substantial financial implications, with annual healthcare costs exceeding $150 billion in the United States alone [1]. The primary causes of nervous system injuries are degenerative disorders and trauma, with peripheral neuropathy affecting 2-7% of the global population [2]. Although peripheral nerves possess regenerative capabilities, the process is slow and often fails to result in full functional recovery even with prompt intervention [3–4].

Peripheral nerve injuries (PNIs) offer various repair options, but little progress has been made in refining treatments over the past few decades. The gold standard approach remains direct repair with or without a nerve graft. Direct repair involves rejoining severed nerve ends, which is suitable for gaps up to 4 mm [5]. Larger gaps necessitate the incorporation of autologous nerves from the patient’s body to repair the damaged site [5–6]. As these methods have considerable limitations, emerging treatments such as localized delivery of exogenous growth factors like nerve growth factor (NGF), glial-derived neurotrophic factor (GDNF), and brain-derived neurotrophic factor (BDNF) have all shown promise, but are quite expensive and their short lifespans in the body can limit their efficaciousness [7–9].

Aside from direct tissue damage and disease [10], the localized production of reactive oxygen species (ROS), defined as secondary neuronal injury [11], hinders nerve regeneration and limits functional outcome recovery [12–13]. Specifically, ROS leads to oxidative damage in nerve fibers, disrupts the blood-nerve barrier, and impairs Schwann cell function that is critical for nerve regeneration [14–18]. Consequently, strategies to reduce ROS levels hold the potential for promoting nerve regeneration and improving functional outcomes as a means of countering ROS-induced neurotoxicity.

An approach for addressing ROS-induced neurotoxicity involves the delivery of small molecules to protect cells and promote neotissue growth. Simple signaling molecules (SSMs) are small bioactive molecules that influence a range of cellular pathways, including neural protection and regeneration. Several SSMs, including glutathione (GSH), n-acetyl cysteine (NAC), hydrogen sulfide (H_2_S), lipoic acid (LA), and carbon monoxide (CO), have been found to exhibit promising neural therapeutic potential, with H_2_S, NAC, and GSH possessing particularly notable benefits which could be leveraged to treat PNIs.

H_2_S, a gaseous signaling molecule produced endogenously from L-cysteine, redistributes GSH to the mitochondria and influences neuronal differentiation and regeneration [19–21]. NAC, a precursor to L-cysteine, is clinically used to counter acetaminophen toxicity that achieves its bioactivity by elevating intracellular GSH levels [22–23]. GSH is an antioxidant tripeptide that maintains two forms - reduced (GSH) and dimerized (GSSG) - whose ratio indicates oxidative stress levels [25–27] and whose elevated levels can result in neuronal regeneration [28]. All of these molecules possess signaling roles within cells and their influence on neural stem cells (NE-4Cs) offers insights into their potential use for the treatment of PNIs through nerve guidance conduits [29–30].

This study assessed H_2_S, NAC, and GSH to determine their optimal therapeutic windows for cytoprotection and neuroinduction to help guide their future incorporation into biomaterials that can achieve desirable controlled payload release. Their ability to counteract oxidative stress was tested using hydrogen peroxide (H_2_O_2_) as a ROS-inducing agent. NE-4Cs were exposed to cytotoxic levels of H_2_O_2_ while concurrently receiving different doses of H_2_S, NAC, and GSH. Cell proliferation, viability, ROS content, and intracellular GSH content were all assessed at varying timepoints. Interestingly, NAC and GSH exhibited enhanced cytoprotective capabilities, while H_2_S facilitated desirable neuronal differentiation behavior.

## MATERIALS & METHODS

### Materials

The experimental materials and reagents were sourced from various suppliers. Sodium hydrosulfide (NaHS) from Sigma Aldrich, N-acetyl cysteine (NAC) from Alfa Aesar, glutathione (GSH and GSSG) from Sigma Aldrich, and hydrogen peroxide (H_2_O_2_) from Ward’s Science. Trition X-100 was obtained from Acros Organics, while 5,5′-dithio-bis(2-nitrobenzoic acid) (DTNB), β-NADPH, glutathione reductase (GR), sulfosalicylic acid, 2-vinylpyradine, and triethanolamine were all sourced from Sigma Aldrich. Additionally, 4% paraformaldehyde was acquired from ThermoFisher for fixation and preservation purposes.

### Cell Culture

NE-4C neural stem cells (ATCC) were cultured in T-75 flasks (Corning) at a density of 300,000 cells per flask using sterile filtered α-MEM media supplemented with 10% FBS and 1% Pen-Strep (Gibco), referred to as α-MEM culture media. Cells were incubated at 37°C with 5% CO_2_, refreshing the media every two days after washing with 1X phosphate buffered saline (PBS). Upon reaching ∼ 80% confluency, cells were passed using 0.05% Trypsin/EDTA. NE-4C for all studies were from passage P5 or greater and plated in 24-well tissue-culture plates (Fisher). Cell used in 3-day studies were plated at 20,000 cells per well, while for 7-day studies, 5,000 cells per well were used, all with a 6-hour adhesion time allotted prior to exposure with experimental media.

Cell culture media was supplemented with a molecule or molecules of interest including the following combinations: NaHS, NAC, GSH, H_2_O_2_, H_2_O_2_ + NaHS, H_2_O_2_ + NAC, and H_2_O_2_ + GSH. A freshly prepared 8 mM NaHS stock solution was used for NaHS-supplemented media. NAC and GSH supplementation employed a 16 mM stock solution, both created by dissolving specific quantities in α-MEM media. These stock solutions underwent sterile filtration (0.22 μm filter) and serial dilution in α-MEM media to yield the concentrations used for experimentation. NaHS concentrations ranged from 3.9 to 500 μM, while NAC and GSH concentrations spanned 0.25 to 16 mM. For H_2_O_2_ supplemented media, a 0.5 M stock solution was diluted accordingly. Additionally, H_2_O_2_ media combined with NaHS, NAC, or GSH was prepared by dissolving these agents into H_2_O_2_-containing media, followed by sterile filtration and serial dilution. Standard α-MEM culture media served as the negative control, with each group plated in quadruplicate (N = 4). Harvesting occurred on days 1, 2, and 3 for 3-day experiments, and on days 1, 3, 5, and 7 for 7-day experiments, with media changes carried out every 2 days.

### Cell Proliferation Assay

Quant-iT PicoGreen dsDNA Assay (ThermoFisher) was employed to quantify cell numbers (N = 4) in each experimental group. Cells were washed with PBS and lysed using 1% Triton X-100 on each experimental day and frozen at - 20 °C, after which samples underwent three freeze/thaw cycles to induce cell lysis. The resulting cell solutions were homogenized using an orbital shaker and sonication. The manufacturer’s assay protocol was followed and fluorescence measurements were taken (ex: 480 nm, em: 520 nm) using a BioTek Cytation 5 spectrophotometer. The fluorescence values were compared to a NE-4C cell standard to determine the final cell count for each sample.

### Cell Viability Assay

ATP levels were measured using the CellTiter-Glo® Luminescent Cell Viability Assay, modifying the manufacturer’s suggested protocol. CellTiter-Glo® reagent was equilibrated to room temperature, added to cellular wells containing α-MEM media, and shaken for 5 minutes to induce cell lysis. The solution was transferred to an opaque 96-well plate, further shaken for 20 minutes, and then analyzed for luminescence (300 - 700 nm) using a BioTek Cytation 5 spectrophotometer. ATP concentration was determined by comparison to readings using a ATP standard curve.

### ROS Detection Assay

Intracellular reactive oxygen species (ROS) levels were measured using the Fluorometric Intracellular ROS Kit, with an adapted protocol for widefield fluorescent microscopy. NE-4C cells were seeded at 20,000 cells per well in α-MEM culture media on coverslips at Day 0. After adhesion, the media was replaced with α-MEM culture media containing various molecules of interest. Following 24 hours of incubation, cells were fixed with 4% paraformaldehyde for 15 minutes, permeabilized with 1% Triton X-100 for 15 minutes, and blocked with 1% bovine serum albumin (BSA) solution in PBS for 30 minutes. Intracellular ROS staining was achieved by treating cells with kit-provided ROS Master Reaction Mix for 1 hour at room temperature. After incubation, cells were washed, mounted on glass slides with ProLong® Diamond Antifade Mountant with DAPI, and imaged using a Zeiss Axiovert 200M inverted widefield fluorescent microscope. Fluorescence measurements were used to visualize cell nuclei (*ex*: 359 nm, *em*: 457 nm) and ROS (*ex*: 520 nm, *em*: 605 nm).

### GSH Assay

GSH and GSSG concentrations were determined by employing an established method [31]. Cells were trypsinized, lysed with extraction buffer (0.1% Triton X-100 and 0.6% sulfosalicylic acid in 0.1 M potassium phosphate buffer at pH 7.4), and frozen. After three thawing cycles were completed, extracts were assessed for GSH and GSSG content. Specifically, GSH was oxidized to TNB with DTNB to determine its level. A companion study was completed where GSSG was first converted to GSH using glutathione reductase (GR) before the DTNB to TNB conversion was conducted. Cyclic TNB production could therefore measure GSH and GSSG (*i.e.*, total minus GSH) content. Fresh DTNB, β-NADPH, and GR solutions were mixed and added to cell extracts. After 30 seconds, β-NADPH was introduced, and the cyclic TNB production rate was measured (*abs*: 412 nm) using a BioTek Cytation 5 spectrophotometer. Total GSH and GSSG content was calculated via GSH standard curve regression.

### Immunofluorescence Staining & Imaging

Wide-field fluorescent microscopy was used to assess the differentiation of NE-4C cells into neurons. At Day 0, coverslips were placed in a 24-well plate, and cells were seeded at 5,000 cells per well in α-MEM media. After similar staining steps outlined for identifying ROS, cells were instead incubated with Alexa Fluor® 647 conjugated β3-Tubulin Antibody. Coverslips were mounted with ProLong® Diamond Antifade Mountant with DAPI, dried for 24 hours, and viewed on a Zeiss Axiovert 200M microscope, for which DAPI (*ex*: 359 nm, *em*: 457 nm) and β3-tubulin (*ex*: 488 nm, *em*: 507 nm) were observed. Using ImageJ, the corrected total cell fluorescence (CTCF) of β3-tubulin was calculated with area, integrated density, and mean intensity values all measured. Randomly selected NE-4C cells (≥ 8) per image were outlined for CTCF calculation, subtracting out mean background fluorescence as determined from multiple dark spots. Mean CTCF per cell was averaged from all cells analyzed, providing supporting information on β3-tubulin expression.

### Statistical Analysis

Data was analyzed using JMP software with Tukey’s Honestly Significant Difference (HSD) test used to determine pairwise statistical differences between groups (p < 0.05). Groups labeled with different letters have a statistically significant variance in their means, while groups that have the same letter, possess statistically insignificant variances in their means.

## RESULTS & DISCUSSION

As SSMs, H_2_S, NAC, and GSH influence endogenous functions making them valuable tools for PNI repair. Their small molecular size and potential therapeutic benefit in cytoprotection and neural regeneration make them ideal to be controllably delivered over time. To establish therapeutic parameters for their release from biomaterials, this research evaluated each molecule for its cytoprotective and inductive potential.

### Simple Signaling Molecule Cytotoxicity

Before exploring the oxidative and neuroinductive effects of H_2_S, NAC, and GSH, a cytotoxic limit for each simple signaling molecule was determined. Neural stem cells (NE-4C) were subjected to different concentrations of the specific SSMs of interest for a duration of up to 7 days. PicoGreen and CellGlo® Luminescence assays were employed to assess cell number and viability, respectively.

### NaHS (H_2_S donor) Cytotoxicity

NE-4C cells were exposed to varying concentrations of H_2_S (3.9 - 500 μM) to identify the upper limit tolerable by cells (**Figure 1**). As H_2_S is gaseous in nature, a donor molecule must be used to deliver it in solution, for which NaHS was chosen. At Day 1, regardless of H_2_S concentration, all NE-4Cs had highly similar cell counts when compared to the negative control cells (*i.e.*, those treated with α-MEM culture media with no H_2_S stimulus, **Figure 1A**). By Day 3, NE-4Cs exposed to 500 μM H_2_S showed limited proliferation which persisted through Days 5 and 7 as compared to the control cells that proliferated considerably over the 7 days of the study. Cells incubated with 125 and 250 μM H_2_S had slightly delayed proliferation compared to the no stimulus control, but nowhere near as severe as the NE-4Cs subjected to 500 μM H_2_S. NE-4Cs exposed to 3.9 - 62.5 μM H_2_S possessed similar growth kinetics as the negative control cells throughout the experiment. These data showed a moderate concentration-based toxicity transition up to the 500 μM NaHS group having a significant, negative impact on stem cell functionality. To complement the proliferation data, ATP concentration was assessed for all stimulus groups at the same time points (**Figure 1B**). ATP concentration across all groups, including 500 μM H_2_S, increased across the days of the experiment. For cells treated with < 500 μM H_2_S, this indicated metabolically active cells where ATP concentration increased as cell count increased. As the cells exposed to 500 μM did not proliferate in the same manner, the elevated ATP concentrations observed in these cells could have been due to cell overdrive and apoptosis rather than healthy metabolic activity [32]. Based on these data, 500 μM H_2_S was determined to be the cytotoxic limit for NE-4Cs and all further experiments did not exceed this concentration.

**Figure 1.**
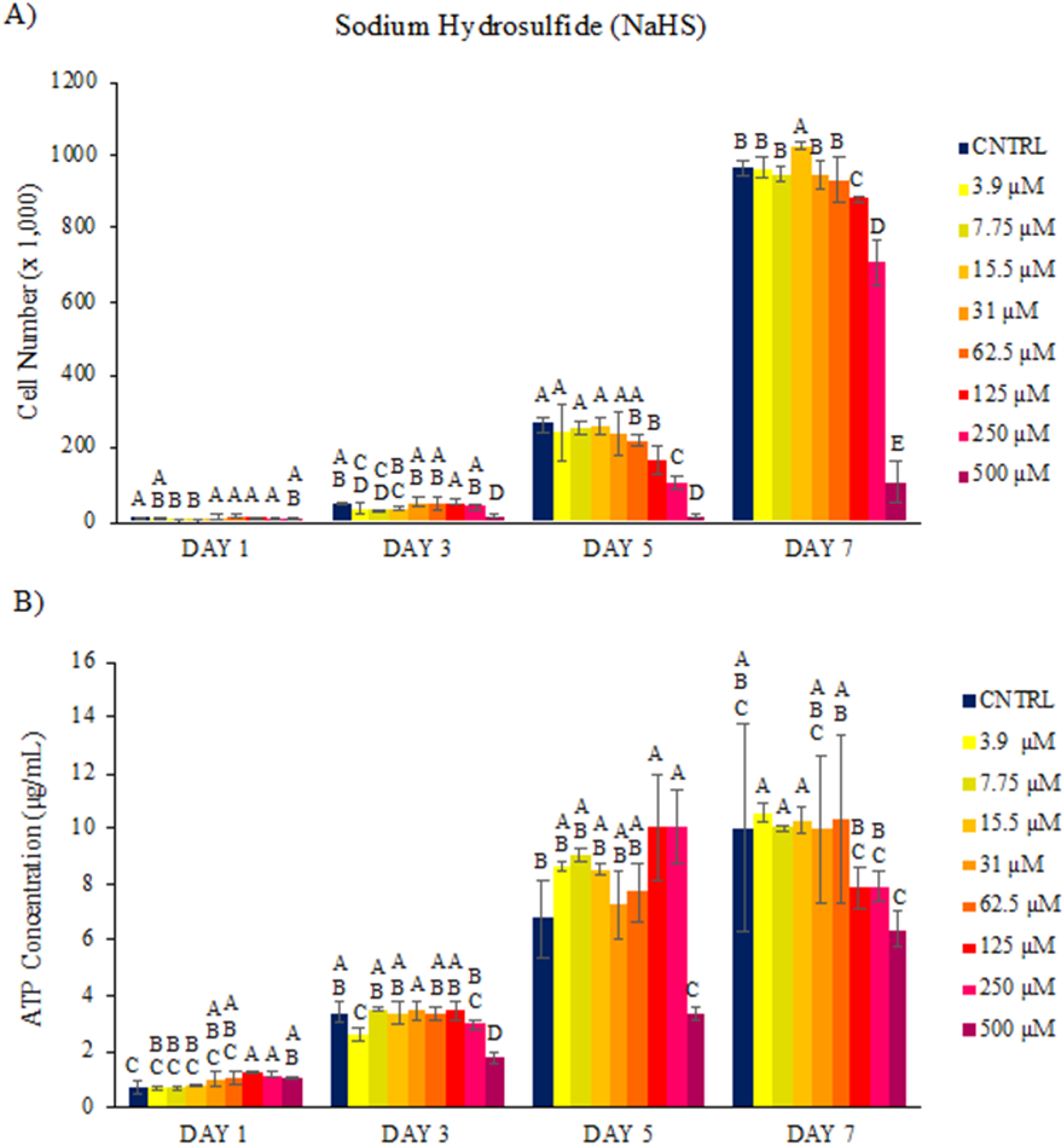
NaHS (H_2_S donor) cytotoxicity in NE-4Cs. The impact of H_2_S from 3.9 - 500 µM was evaluated over 7 days. Through NE-4C cell proliferation data (A) and ATP concentration data (B), a cytotoxic limit of 500 µM H_2_S was determined. Groups that possess different letters have statistically significant differences (p < 0.05) in mean whereas those that possess the same letter are statistically similar (N = 4).

### NAC Cytotoxicity

NE-4C cells were subjected to different NAC concentrations (0.25 - 16 mM) to establish its cytotoxic limit for these cells (**Figure 2**). It is important to note that these concentrations are on the mM scale, far greater than the μM scale used for H_2_S as NAC is a less potent molecule. At Day 1, regardless of NAC concentration, all NE-4Cs had similar or greater cell numbers when compared to the negative control cells (*i.e.*, those treated with α-MEM culture media with no NAC stimulus, **Figure 2A**). By Day 3, cells exposed to 1 or 2 mM NAC exhibited slightly elevated cell numbers in comparison to the control group suggesting this molecule may be mildly mitogenic. That being said, by Days 5 and 7, the negative control cells and NE-4Cs incubated with 0.25, 0.5, or 4 mM NAC caught up to the cell counts observed for the 1 and 2 mM NAC experimental groups. For NE-4Cs exposed to 8 mM NAC, there was a drop in cell population from Day 5 to Day 7. This could be due to a delayed cytotoxic effect felt by the cells after being subjected to NAC for more than 5 days. Cells cultured in the highest concentration of NAC (i.e., 16 mM) never proliferated, remaining right around the starting cell count of 5,000 cells for the entire 7 days of the study.

**Figure 2.**
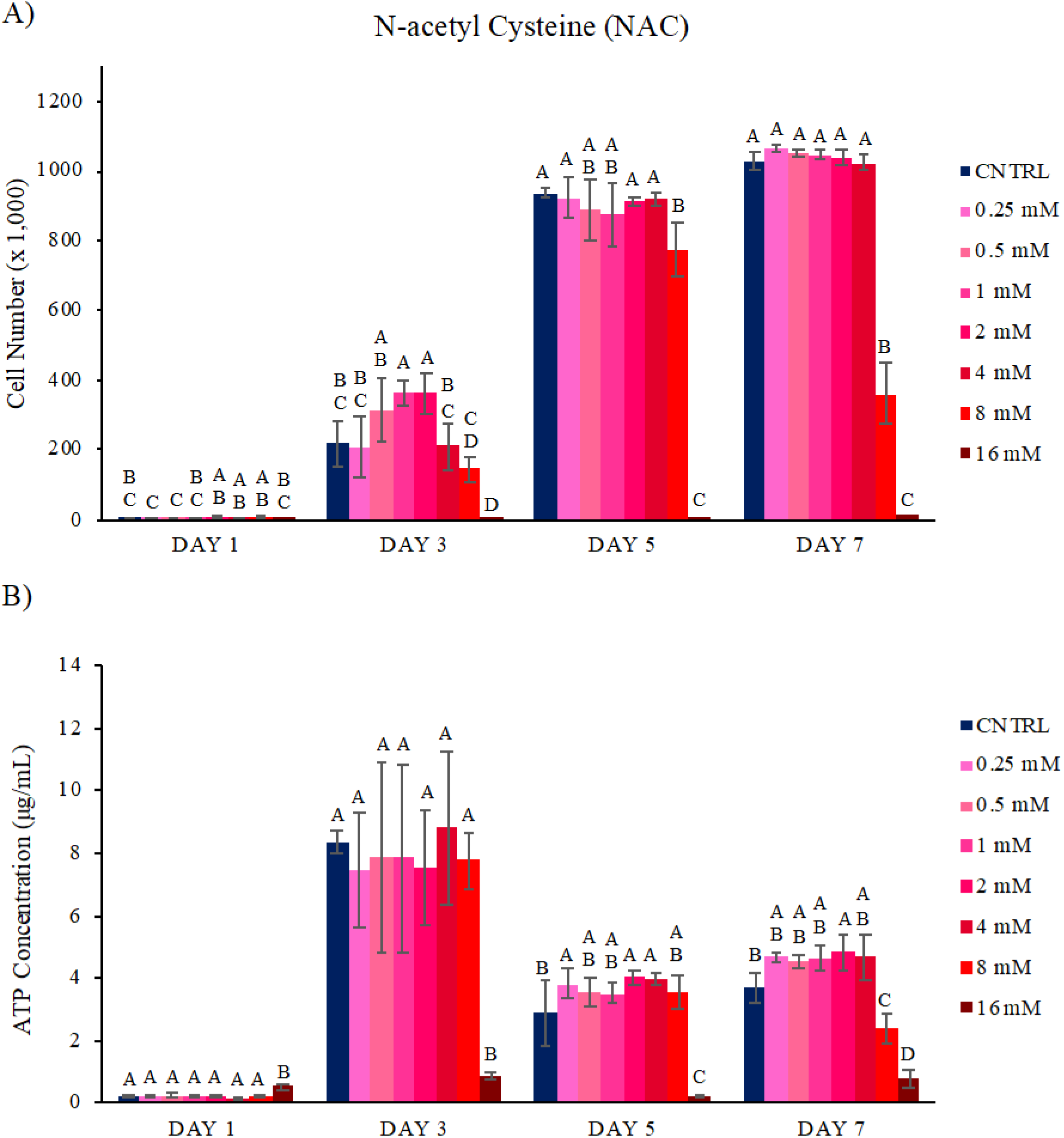
NAC cytotoxicity in NE-4Cs. The impact of NAC from 0.25 - 16 mM was assessed over 7 days. Through NE-4C cell proliferation data (A) and ATP concentration data (B), a cytotoxic limit of 16 mM NAC was calculated. Groups that possess different letters have statistically significant differences (p < 0.05) in mean whereas those that possess the same letter are statistically similar (N = 4).

To supplement the proliferation data, ATP concentration was evaluated for all stimulus groups at the same time points (**Figure 2B**). For NE-4Cs subjected to 0.25 - 4 mM NAC, ATP concentration was similar to the negative control cells with little metabolic activity on Day 1, peaking on Day 3, and moderate levels on Days 5 and 7. This behavior is consistent with the proliferation data that showed confluency was reached by Day 5, in which the high metabolic activity associated with cell mitosis would have been minimized [33]. Cells incubated in 8 mM NAC exhibited similar metabolic behavior as their proliferative profile with the highest ATP concentration observed on Day 3 when they were amply growing, moderate levels at Day 5 as they reached confluency, and lower levels at Day 7 as their cell population decreased. For NE-4Cs cultured in 16 mM NAC, limited metabolic activity was observed over 7 days aligning with the non-proliferative behavior of these cells. These results indicate that 16 mM NAC is cytotoxic to NE-4C cells, so all experiments to follow did not surpass this concentration.

### GSH Cytotoxicity

NE-4C cells were incubated in escalating GSH concentrations (0.25 - 16 mM) to determine the maximum amount these cells can endure (**Figure 3**). This range is consistent with the one used for NAC and much greater than that of H_2_S. At Day 1, all NE-4Cs exhibited similar cell proliferation as the negative control cells (*i.e.*, those treated with α-MEM culture media with no GSH stimulus, **Figure 3A**). At Day 3, the cells subjected to 0.25 - 8 mM GSH were in similar or significantly greater number than the negative control, demonstrating a potentially mitogenic bioactivity of GSH though this effect did not appear to be concentration dependent. That being said, by Days 5 and 7, the negative control cells and NE-4Cs incubated with 0.25 - 2 mM NAC all had similar cell counts. Cells exposed to 4 mM or 8 mM NAC reached a plateau cell number about 60% of those exposed to no NAC or lower concentrations of NAC. For the 16 mM GSH treated cells, no cell proliferation occurred over the 7 days of the study. To further support the proliferation data, ATP concentration was evaluated for all stimulus groups at the same time points (**Figure 3B**). For cells exposed to 0.25 - 2 mM GSH, ATP was greatest at Day 3 corresponding with cell confluency being achieved by Day 5 and its influence on metabolic activity being observed. The cells incubated with 4 mM or 8 mM GSH had less ATP at Day 3 then the 0.25 - 2 mM GSH treated cells, which seemed to level off through Days 5 and 7. This paralleled the proliferation data where these cells did not proliferate significantly past Day 3. These observations could indicate a variance in toxicity, where some cells are still dividing while others are being affected by these GSH concentrations, resulting in a relatively static cell number. The elevated levels of ATP at Day 5 could then be suggestive of cellular apoptosis within this population [35]. Considering this information, it was determined that 16 mM GSH was the cytotoxic limit to be used for NE-4C cell studies going forward.

**Figure 3.**
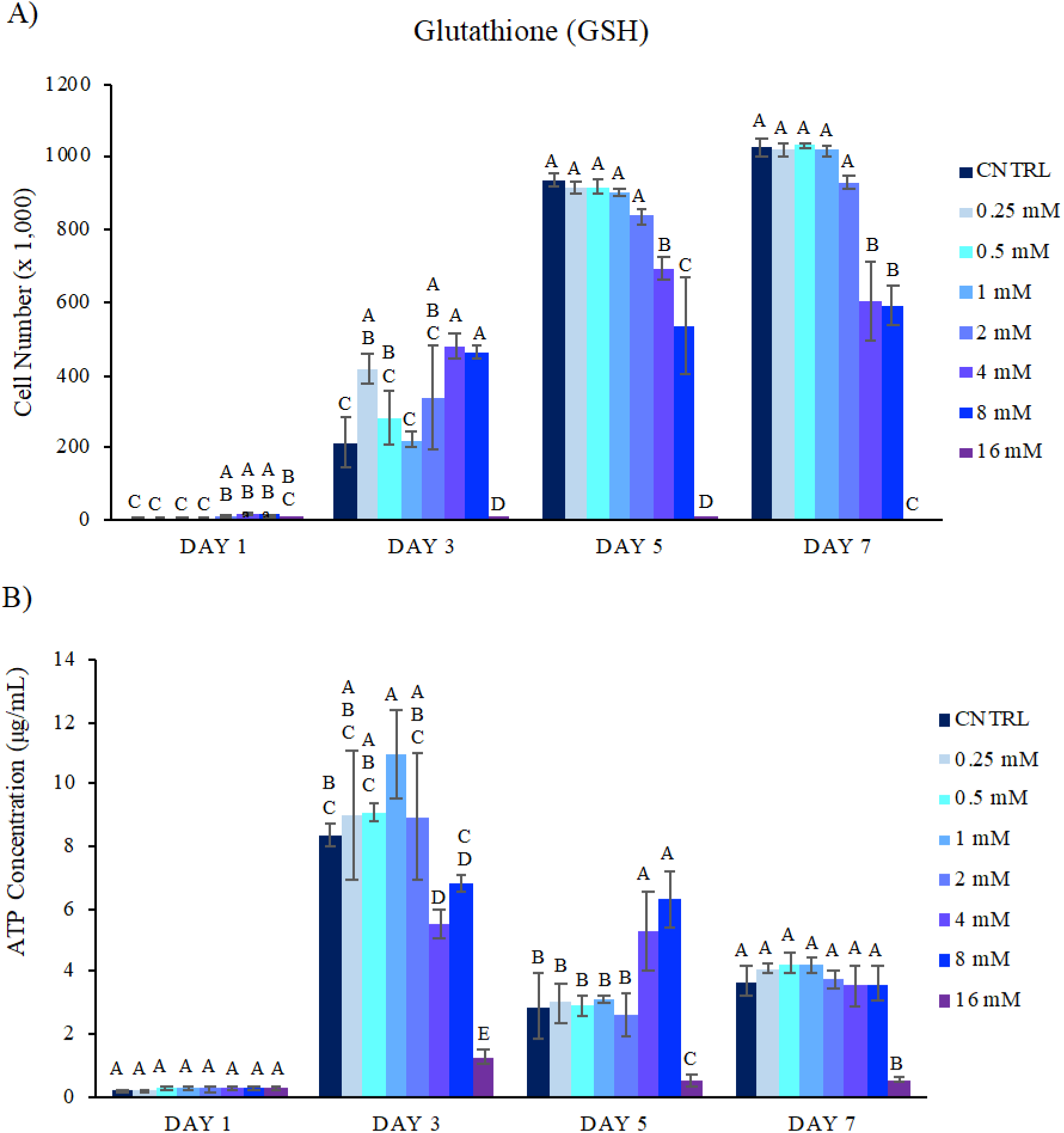
GSH cytotoxicity in NE-4Cs. The impact of GSH from 0.25 - 16 mM was studied over 7 days. Through NE-4C cell proliferation data (**A**) and ATP concentration data (**B**), a cytotoxic limit of 16 mM NAC was measured. Groups that possess different letters have statistically significant differences (p < 0.05) in mean whereas those that possess the same letter are statistically similar (N = 4).

### Oxidative Stress Evaluation

To study the capacity H_2_S, NAC, and GSH have to protect NE-4Cs against oxidative stress, hydrogen peroxide (H_2_O_2_) was utilized as a stressor. H_2_O_2_ was chosen due to its toxic nature as a known ROS within the body and its vast history of use in oxidative stress models [24, 33–35]. To help determine the concentration of H_2_O_2_ that causes oxidative stress in NE-4Cs, a cytotoxic screen of H_2_O_2_ was conducted over two days with a starting cell number of 20,000 cells per well (**Figure 4**). After just 1 day, any concentration greater than or equal to 150 μM H_2_O_2_ appeared to be cytotoxic to the cells. These NE-4Cs had significantly fewer cells than the control group as well as lower cell counts than the starting cell number (*i.e.*, 20,000). These cells also did not recover any proliferative capacity by Day 2, so 150 μM H_2_O_2_ was utilized for all follow-on oxidative stress experiments. As oxidative stress and inflammation occur rapidly following an injury [26, 36], studies examining the cytoprotection of various SSMs were conducted over 3 days.

**Figure 4.**
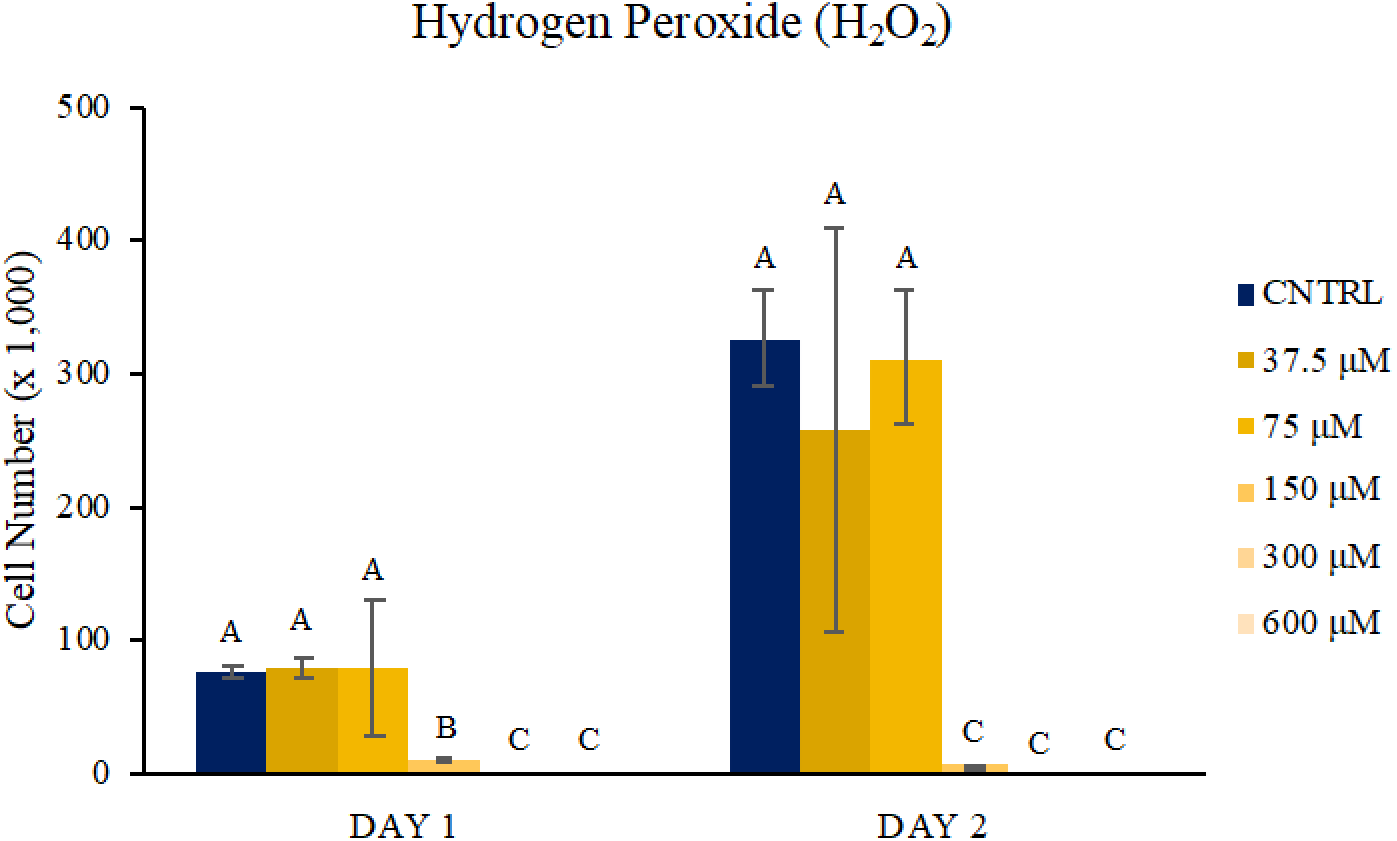
H_2_O_2_ cytotoxicity in NE-4Cs. The impact H_2_O_2_ had on NE-4C cell proliferation was evaluated over 2 days. A cytotoxic limit of 150 μM H_2_O_2_ was calculated. Groups that possess different letters have statistically significant differences (p < 0.05) in mean whereas those that possess the same letter are statistically similar (N = 4).

### H_2_S, NAC, and GSH Supplemented, H_2_O_2_-Treated Cell Cytotoxicity

To investigate the mitigating impact H_2_S, NAC, and GSH have on H_2_O_2_ cytotoxicity, the PicoGreen assay was employed to assess cell proliferation. When positive cell growth kinetics were observed, the CellGlo® Luminescence Assay was also employed to analyze cellular metabolic activity. Once NE-4Cs had adhered to the plate, they were treated with mixed media containing 150 μM H_2_O_2_ supplemented with varying concentrations of H_2_S, NAC, or GSH. Negative and positive controls consisted of α-MEM culture media with no added stimulus and 150 μM H_2_O_2_ alone in α-MEM culture media, respectively. SSM-based cytoprotection for H_2_O_2_-stressed NE-4Cs was considered to be achieved when cell counts and ATP concentration were at least 80% that of cells incubated with α-MEM culture media alone after three days of treatment.

### H_2_S Oxidative Stress Rescue

To examine the cytoprotective effects of H_2_S, NE-4Cs were subjected to 150 μM H_2_O_2_ media complemented with differing concentrations of NaHS (H_2_S donor, 7.75 - 500 μM, **Figure 5**). After 1 day, none of the H_2_S concentrations had a significant H_2_O_2_ mitigating effect. Specifically, all the cells exposed to 150 μM H_2_O_2_, regardless of how much H_2_S was present, had comparable cell counts as the H_2_O_2_ control cells, while the α-MEM culture media control cells proliferated significantly. By Day 3, there still was essentially no cell growth with any of the experimental treatments, so the ATP analysis was not carried out for this study. This indicated that with this model, H_2_S, as delivered via NaHS, did not appear to have a cytoprotective effect for NE-4Cs against the H_2_O_2_ insult. This result was somewhat unexpected as the majority of literature exploring the impact of ROS inducers (H_2_O_2_, NaN_3_, CoCl_2_, oxygen-glucose deprivation/reoxygenation, *etc.*) on neural-related cells *in vitro* employing H_2_S delivery (NaSH or H_2_S-releasing molecule) showed a cytoprotective effect [37–41]. The most likely reason for this difference is that these studies focused on pretreating cells with H_2_S prior to ROS inducer exposure as compared to the H_2_S/H_2_O_2_ co-exposure used in this work. Also, in most cases, stem cells exhibit heightened sensitivity to the detrimental effects of ROS when compared to their differentiated cell counterparts [42–43]. This has been specifically shown with *in vitro* neural injury models comparing unmodified and differentiated neural stem cells (PC12) [44–45]. As this characterization has not been done on undifferentiated NE-4Cs, a similar lack of H_2_S-mediated recovery from H_2_O_2_-induced cell injury was not completely surprising.

**Figure 5.**
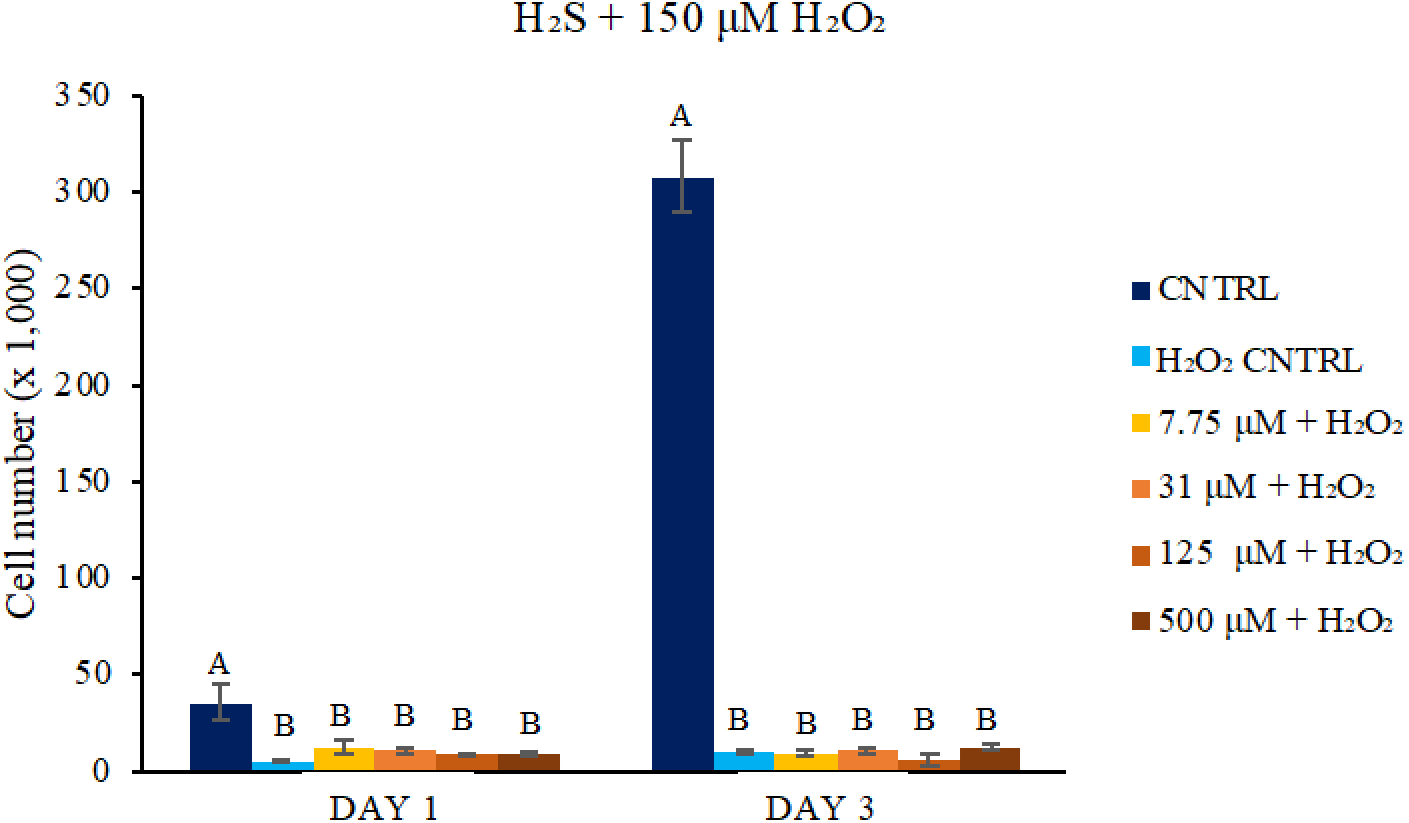
H_2_S influence on H_2_O_2_ cytotoxicity in NE-4Cs. The impact 7.75 - 500 µM H_2_S had on mitigating oxidative stress was explored over 3 days. Through NE-4C cell proliferation data, no significant H_2_S-mediated cytoprotective effect was observed. Groups that possess different letters have statistically significant differences (p < 0.05) in mean whereas those that possess the same letter are statistically similar (N = 4).

### NAC Oxidative Stress Rescue

NE-4Cs were cultured with 150 μM H_2_O_2_ supplemented with varying concentrations of NAC (0.25 - 16 mM, **Figure 6**) to study the cytoprotective capabilities of NAC. After 1 day, H_2_O_2_-agitated cells treated with 1 - 16 mM NAC all had significantly greater cell numbers than the H_2_O_2_ only control group, indicating initial cytoprotection from oxidative stress was mediated by NAC (**Figure 6A**). Although the cell counts within these concentrations were not as substantial as the negative control cells, a similar trend was observed at Day 2, where cells subjected to 1 - 12 mM NAC continued to proliferate. In contrast, cells exposed to the H_2_O_2_ control did not recover their initial starting cell number of 20,000 cells, and the lowest (*i.e.*, 0.25 and 0.5 mM) and highest (*i.e.*, 16 mM) concentrations of NAC also maintained comparably low cell numbers. By Day 3, the H_2_O_2_ shocked cells incubated with 1 - 8 mM NAC expressed similar cell numbers to those cultured in the α-MEM culture media alone control. Also, while the 0.5 mM and 12 mM NAC treated cells had not proliferated as much by this time point, they were still in much greater number than the positive control. The lowest dose (*i.e.*, 0.25 mM) and highest dose (*i.e.*, 16 mM) of NAC showed no impact in mitigating H_2_O_2_ cell cytotoxicity throughout the study. As cells cultured with 1 - 8 mM NAC had their proliferative capacity restored to at least 80% of their normative level, this concentration range is suggested to be near this SSM’s cytoprotective therapeutic window. To investigate these findings further, the CellGlo® Luminescence Assay was then employed to analyze metabolic activity. ATP concentrations measured with the CellGlo® Luminescence Assay detailing the cytoprotective effect of NAC corresponded well with the PicoGreen cell proliferation data (**Figure 6B**). Elevated ATP concentrations, in contrast to the H_2_O_2_ treated control cells were observed in the NE-4Cs treated with 2 - 12 mM NAC and H_2_O_2_ throughout the entire three days of the study, with ATP leveling off between Day 2 and Day 3 as the cells began to reach confluency. These data combined with the cell proliferation data, not only indicated the impact NAC can have on cell survival but also on keeping those cells metabolically active. It is interesting to note, that although there was not much cell proliferation in the H_2_O_2_ only, 0.25 mM NAC + H_2_O_2_, or 16 mM NAC + H_2_O_2_ treated cells, these cells still appeared to have higher ATP levels despite their low cell number. Elevated concentrations of ATP can be associated with the high energy requirement of cellular apoptosis, so this trend is more likely an indicator of cell death rather than healthy metabolic activity [32]. These ATP observations helped to narrow the cytoprotective window of NAC against H_2_O_2_-medicated oxidative stress to 2 - 8 mM.

**Figure 6.**
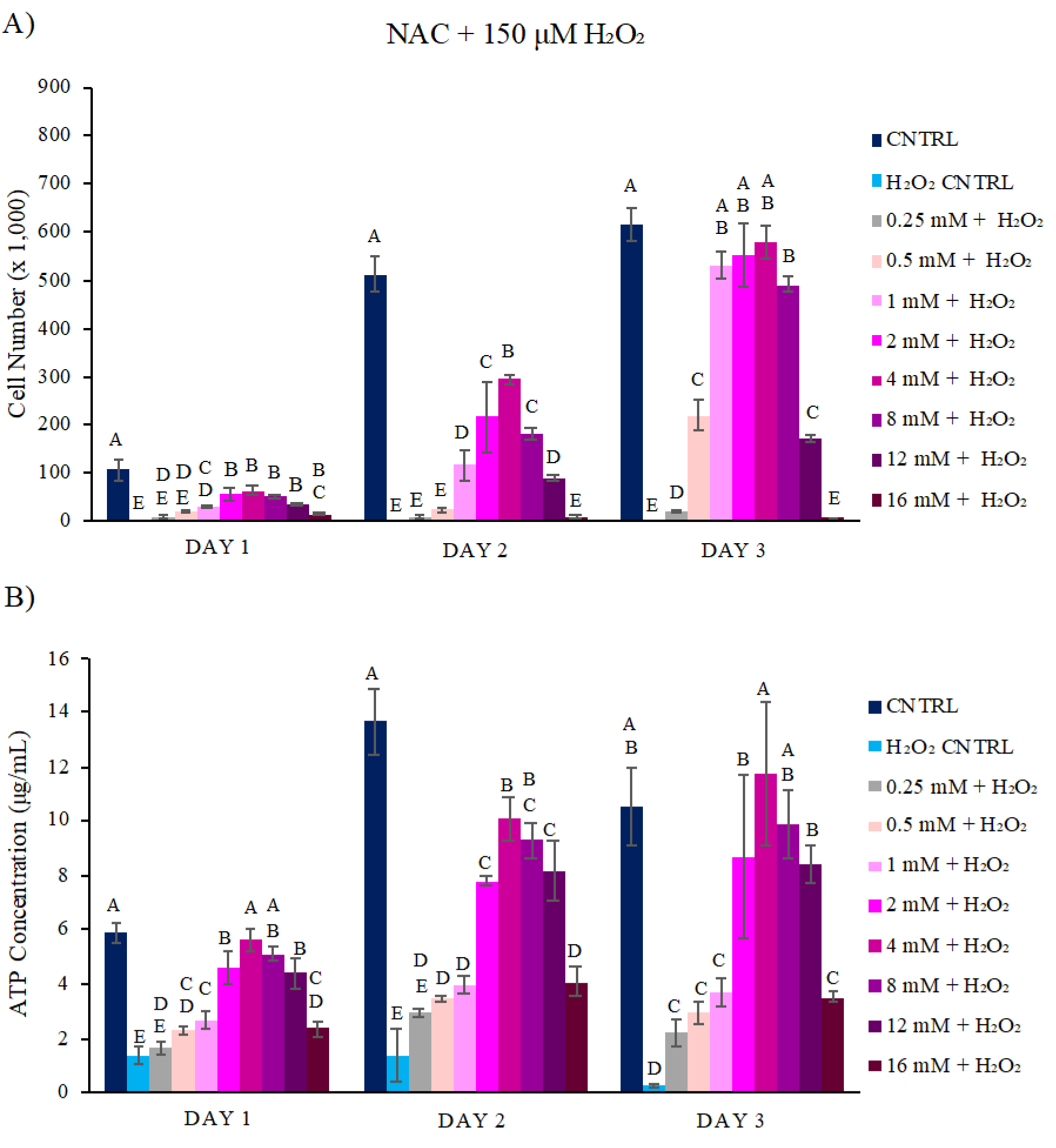
NAC influence on H_2_O_2_ cytotoxicity in NE-4Cs. The impact 0.25 - 16 mM NAC had on mitigating oxidative stress was studied over 3 days. Through NE-4C cell proliferation data (**A**) and ATP concentration data (**B**), a cytoprotective window of 2 - 8 mM NAC was uncovered. Groups that possess different letters have statistically significant differences (p < 0.05) in mean whereas those that possess the same letter are statistically similar (N = 4).

### GSH Oxidative Stress Rescue

NE-4Cs were exposed to 150 μM H_2_O_2_ mixed with a range of GSH concentrations (0.25 - 16 mM, **Figure 7**) to investigate the cytoprotective effects of this SSM. After 1 day, NE-4Cs exposed to the adverse H_2_O_2_ stimulus with 0.5 - 16 mM GSH exhibited greater cell proliferation than the H_2_O_2_ control group and increased cell numbers in a mostly comparable manner to that of the negative control (**Figure 7A**). By Day 2, similar overall growth trends were observed for stressed cells subjected to 1 - 12 mM GSH. In contrast, cells exposed to 0.5 or 16 mM GSH showed minimal proliferative effects though all these concentrations (*i.e.*, 0.5 - 16 mM GSH) had significantly higher cell numbers than the H_2_O_2_-exposed control cells. By Day 3, NE-4Cs exposed to 0.5 - 1 mM GSH in addition to H_2_O_2_ had proliferated some but did not have as high of cell counts as those treated with 2 - 12 mM GSH. Cells incubated with 150 μM H_2_O_2_ and 0.25 mM GSH showed limited proliferative capacity similar to that of NE-4Cs provided H_2_O_2_ alone.

**Figure 7.**
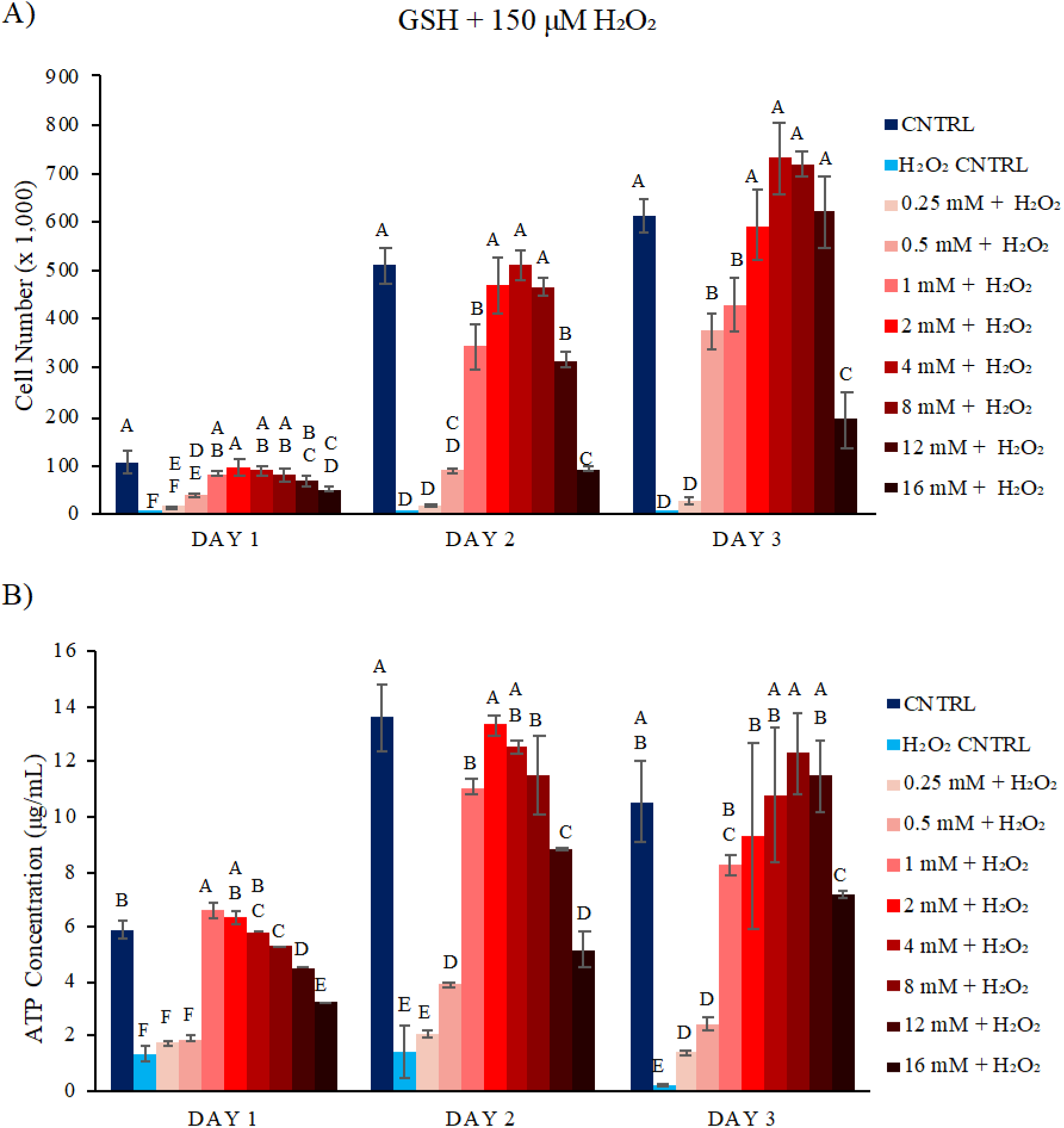
GSH influence on H_2_O_2_ cytotoxicity in NE-4Cs. The impact 0.25 - 16 mM GSH had on mitigating oxidative stress was investigated over 3 days. Through NE-4C cell proliferation data (**A**) and ATP concentration data (**B**), a cytoprotective window of 2 - 12 mM GSH was determined. Groups that possess different letters have statistically significant differences (p < 0.05) in mean whereas those that possess the same letter are statistically similar (N = 4).

Interestingly, 16 mM GSH given without H_2_O_2_ was found to be quite toxic to NE-4Cs (**Figure 3A**), however, when combined with 150 μM H_2_O_2,_ cell counts increased by nearly an order of magnitude over their initial cell count (*i.e.*, 20,000) after three days. This could indicate some extracellular interactions between GSH and H_2_O_2_ that would influence the quantity of both molecules in solution[47]. Based on these results, a range of 2 - 12 mM GSH appeared to have cytoprotective effects against H_2_O_2_ for NE-4C cell proliferation, although 0.5 and 1 mM GSH still restored cell populations to ∼ 60% of what was found with the negative control. To further examine these findings, the CellGlo® Luminescence Assay was employed to measure the metabolic activity of the NE-4Cs used in this model.

The ATP concentration of the cell populations stressed with H_2_O_2_ and co-incubated with GSH (**Figure 7B**) indicated a similar trend as the cell proliferation data. At Days 1, 2, and 3, cells exposed to 150 μM H_2_O_2_ supplemented with 1 - 12 mM GSH exhibited elevated ATP concentrations in contrast to the H_2_O_2_ control. By Day 3, this concentration window also induced cells to have similar levels of ATP to those treated with α-MEM culture media alone. H_2_O_2_-agitated cells subjected to 0.25 mM or 0.5 mM GSH maintained the same, relatively low ATP values throughout the three days of the study. Just as there was cell proliferation in the cells exposed to H_2_O_2_ combined with 16 mM GSH, there also was greater ATP concentration, further highlighting the possibility GSH and H_2_O_2_ interacted directly. As H_2_O_2_-stressed NE-4Cs treated with 2 - 12 mM GSH demonstrated both cell proliferation and ATP production greater than 80% of the negative control, it was concluded that these concentrations of GSH have a cytoprotective effect mitigating oxidative stress in this model.

### NAC and GSH Supplemented, H_2_O_2_-Treated Cell Intracellular Evaluation

Though the cytoprotective capabilities of NAC and GSH were valuable findings, the potential for NAC and GSH to be directly oxidized by H_2_O_2_ [47] leaves the possibility that these effects occurred extracellularly, intracellularly, or a combination thereof. To further probe this, intracellular reactive oxygen species (ROS) content was investigated for both NAC and GSH treated H_2_O_2_-stressed cells. Additionally, as GSH is the main intracellular antioxidant [25], its total intracellular content (*i.e.*, the combination of monomeric GSH and dimeric glutathione disulfide - GSSG) was quantified to determine whether NAC influenced the concentrations and relative ratio of GSH and GSSG. Data from GSH-treated cells was not included as the exogenous treatment of GSH acts as a confounding variable for this experiment. As there are greater quantities of NAC and GSH than H_2_O_2_ (*i.e.*, 0.25 - 16 mM versus 150 μM), the intracellular presence of ROS would indicate that not all the H_2_O_2_ was consumed by the SSM extracellularly. Also, if an elevation in total intracellular GSH content (*i.e.*, GSH + GSSG) comparable to the negative control was found, this would suggest a cytoprotective pathway for NAC [25].

### ROS Presence

To assess intracellular ROS levels in the observed therapeutic ranges for NAC or GSH, cells were subjected to α-MEM culture media alone (negative control) or supplemented with 150 μM H_2_O_2_ with 0 mM (positive control), 0.25 mM, 1 mM, 4 mM, or 16 mM NAC or GSH. After 1 day of incubation, cells were fixed with 4% paraformaldehyde and assayed using the Fluorometric Intracellular ROS Detection Assay supplemented with DAPI staining (**Figure 8**). For all stimuli used, the images show the cell nuclei (DAPI, *ex:* 359 nm, *em:* 457 nm) and ROS (fluorescent detection agent, *ex:* 520 nm, *em:* 605 nm). Each image is representative of triplicate samples for each treatment group.

**Figure 8.**
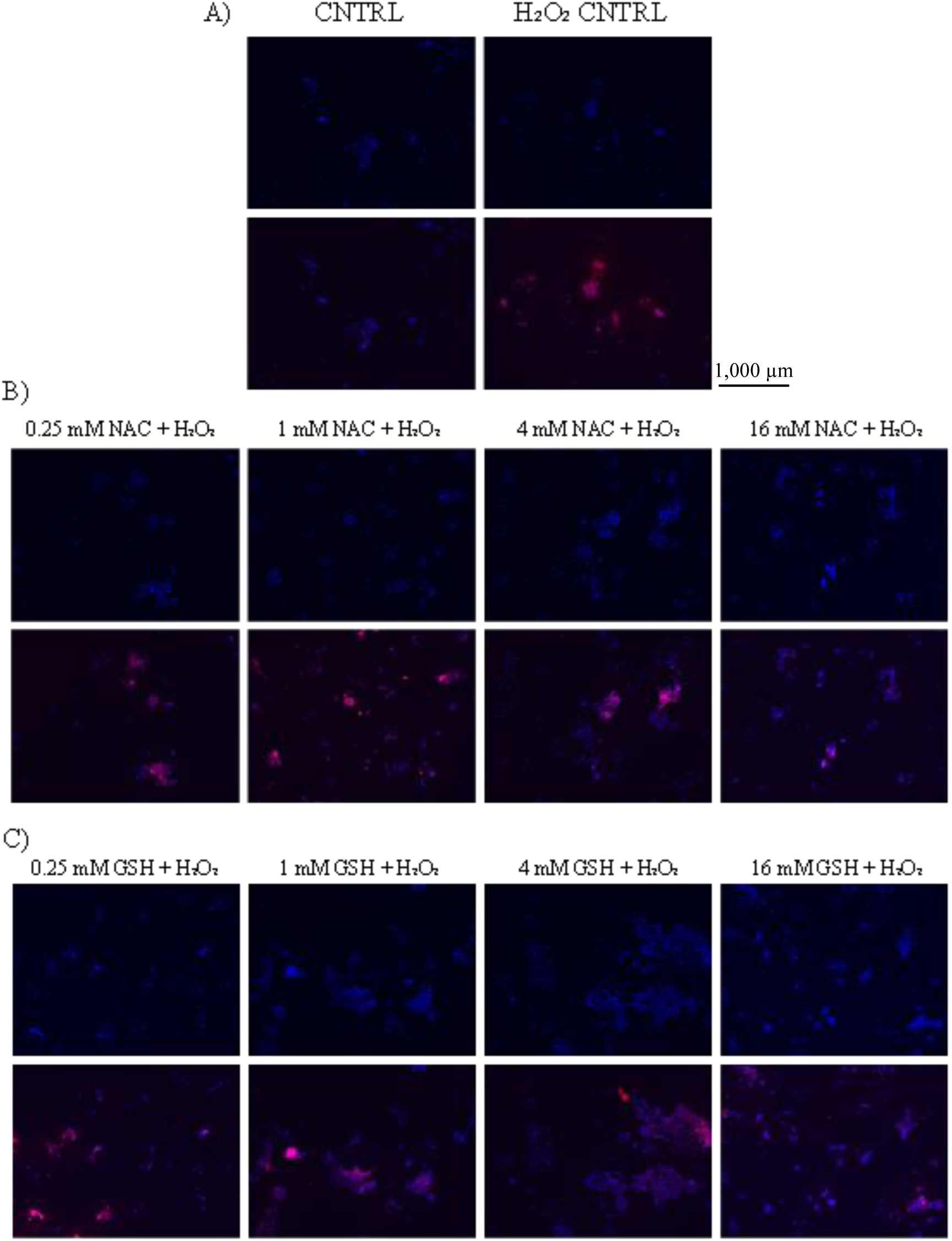
NAC and GSH impact on intracellular ROS presence. Media with and without 150 μM H_2_O_2_ (**A**), NAC + 150 μM H_2_O_2_ (**B**), or GSH + 150 μM H_2_O_2_ (**C**) was used to treat cells for which their nucleus and intracellular ROS were stained. DAPI images alone are displayed in the top panel and DAPI and ROS overlay images are shown in the bottom panel for each group (N = 3).

**Figure 9.**
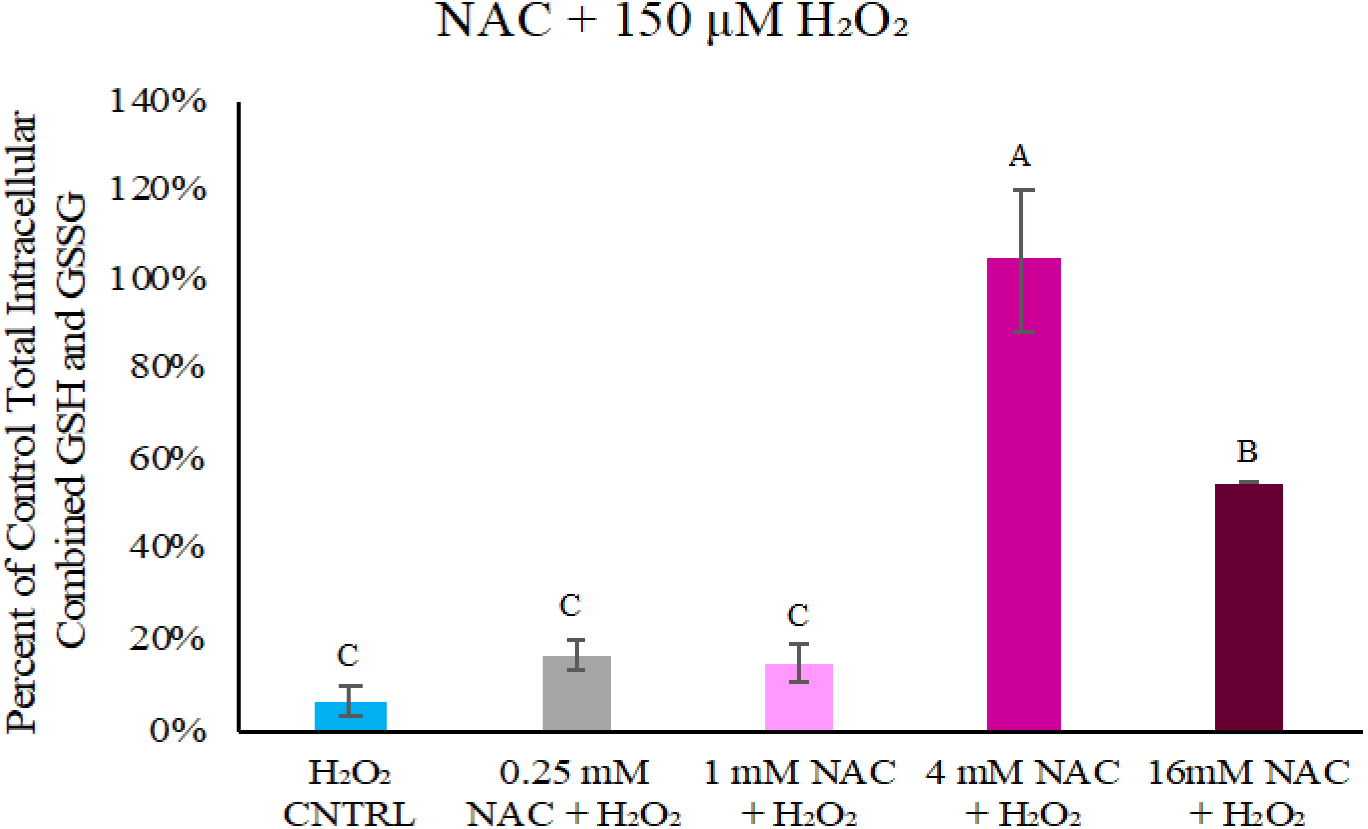
NAC impact on intracellular GSH content in NE-4Cs. The influence 0.25 - 16 mM NAC had on altering GSH content from 150 µM H_2_O_2_ agitated cells was studied over 1 day. These data are presented as a percent of healthy control cells solely subjected to α-MEM culture media with 4 mM NAC showing a complete return of normative total intracellular GSH levels. Groups that possess different letters have statistically significant differences (p < 0.05) in mean whereas those that possess the same letter are statistically similar (N = 3).

The negative and positive controls showed ROS presence is highly stimulus dependent. Specifically, very little ROS fluorescence was observed in NE-4Cs cultured in α-MEM culture media alone whereas nearly every cell exposed to the 150 μM H_2_O_2_ insult fluoresced, showing the intracellular presence of ROS (**Figure 8A**). For stressed cells cultured with 0.25 - 16 mM NAC, ROS-labeled fluorescence was observed across all treatment groups (**Figure 8B**). Those given the lower NAC doses (*i.e*., 0.25 and 1 mM) possessed similar ROS levels to that of the 150 μM H_2_O_2_ control. This observation aligned with the cell cytotoxicity data, as the cell number and metabolic activity were similarly reduced at these same concentrations after 1 day (**Figure 6**).

For H_2_O_2_-stressed cells incubated with 4 mM NAC, ROS expression was still observed, though not in every cell like was seen with 0.25 and 1 mM NAC. This result suggests that a therapeutic concentration of this SSM may be mitigating the ROS content within NE-4Cs. Interestingly, 16 mM NAC yielded the lowest ROS presence though this is likely due to its inherent cytotoxicity (**Figure 4**) as well as the cell death seen when co-delivered with H_2_O_2_ (**Figure 6**). Taken together, these findings complement the earlier work establishing a NAC therapeutic window of 2 - 8 mM.

For NE-4Cs incubated with H_2_O_2_ and GSH, ROS-associated fluorescence was seen throughout the entire range of GSH concentrations examined (**Figure 8C**). Almost all stressed cells exposed to 0.25 or 1 mM GSH possessed intracellular ROS. For the lowest concentration, this result aligns with the cell growth suppression and diminished ATP content found with these stimuli 1-day post-exposure (**Figure 7**). Interestingly, for 1 mM GSH, the considerable presence of ROS did not impact cell function and health at Day 1 though this could play a role in the non-confluency cell count plateau seen for these cells at Day 3 (**Figure 7A**) [46]. For cells subjected to 4 mM GSH and 150 μM H_2_O_2_, ROS-based fluorescence was observed, but not at the same intensity as the H_2_O_2_ control or lower GSH concentrations. This result is consistent with the cytoprotective effect seen with this GSH concentration (**Figure 7**). Cells incubated in 16 mM GSH with an adverse stimulus (*i.e.*, H_2_O_2_) expressed the least amount of ROS in comparison to the other concentrations of GSH. NE-4C cytotoxicity was previously noted with 16 mM GSH with and without H_2_O_2_ supplementation in **Figure 3** and **Figure 7**, respectively, when compared to cells cultured in α-MEM culture media alone. These data align with the previous results determining a GSH therapeutic window of 2 - 12 mM.

When considering this data in aggregate, it suggests that cell-mediated effects are primarily responsible for the influence NAC and GSH have on mitigating H_2_O_2_-induced oxidative stress. The considerable and similar ROS content seen within cells receiving either SSM at the lower two concentrations tested (*i.e.*, 0.25 and 1 mM) provide supports that the H_2_O_2_ is not undergoing decomposition even though there is 1.67 - 6.67 times as much SSM present. In addition, the antioxidant kinetics of NAC and similar thiols like GSH and cysteine have notably slower reaction rates (0.16, 0.89, and 2.9 M^−1^ s^−1^, respectively) than antioxidant enzymes (peroxiredoxins - 1x10^7^ - 4x10^7^ M^−1^ s^−1^ and superoxide dismutase - 2.3x10^9^ M^−1^ s^−1^) [47]. This is mainly due to physiological thiolate concentrations derived from the pKa values of these thiols: NAC < GSH < cysteine [48]. Consequently, the reactivity of NAC with reactive oxygen species like O_2_^-^, H_2_O_2_, and ONOO^-^ is relatively restrained and less physiologically significant [49]. However, extracellular NAC can potentially lead to intracellular conversion to cysteine, contributing to pathways involved in GSH synthesis, and sulfane sulfur species directly or indirectly through H_2_S production [50].

At the highest SSM concentration assessed (*i.e.*, 16 mM), less intracellular ROS was present, yet the proliferative and ATP content were not fully rescued indicating the cytotoxicity of these molecules at this concentration governed the lack of healthy cells in culture. Additionally, these results were unable to determine whether NAC or GSH is more effective at decreasing intracellular ROS. Instead, the data suggests that either SSM has desirable effects provided they are delivered at concentrations within their therapeutic window.

### Intracellular GSH Assessment

Total intracellular GSH (*i.e.*, GSH+GSSG) was assayed to further investigate the capacity NAC has to modulate H_2_O_2_-induced oxidative stress in NE-4Cs. The exogenous treatment of GSH itself could affect the validity of this assay, therefore the assay was only conducted with NAC-treated cells. As GSH is the major antioxidant within cells, this measure is valuable in assessing whether NAC influenced these levels within a stressful environment [23, 51]. Cells were subjected to α-MEM culture media alone (negative control) or supplemented with 150 μM H_2_O_2_ with 0 mM (positive control), 0.25 mM, 1 mM, 4 mM, or 16 mM NAC. After 1 day of incubation, cells were harvested and analyzed. These data are presented as a percent of the total intracellular GSH for the healthy control cells incubated with α-MEM culture media alone (**Figure 8**).

For the H_2_O_2_ control cells, total intracellular GSH was diminished to less than 10% of the healthy negative control cells solely incubated with α-MEM culture media (**Figure 8**). For the cells exposed to 0.25 mM or 1 mM NAC with 150 μM H_2_O_2_, there was not a statistically significant increase in total intracellular GSH when compared to the H_2_O_2_-exposed positive control cells. This corresponded well with the ROS fluorescence data (**Figure 8B**) where high intracellular ROS content was observed, as well as the cytotoxicity data **(Figure 6**) where these cells had a low cell number after 1 day. This supports the location of the low end of the therapeutic window, where 0.25 - 1 mM NAC may be insufficient in protecting NE-4Cs from the oxidative stress caused by H_2_O_2_. For the H_2_O_2_-stressed cells subjected to 4 mM NAC, a concentration within the identified therapeutic window, total intracellular GSH was similar to the negative control cells in α-MEM culture media and significantly greater than the H_2_O_2_-treated positive control cells. The ROS-based fluorescence for these cells was less than the H_2_O_2_ control cells and greater than the α-MEM culture media control cells (**Figure 8A & B**). Although there was still ROS present in the cells exposed to 4 mM NAC, there was similar amounts of total intracellular GSH as the healthy control cells. This valuable information indicates that NAC may be playing a role in elevating GSH levels as a cytoprotective measure against the oxidative insult. The stressed cells treated with 16 mM NAC also had an elevated total intracellular GSH levels when compared to the H_2_O_2_ control cells but only ∼ 50% of the content found within the negative control cells. This also could be representative of the effect NAC has on total intracellular GSH as there was little ROS fluorescence shown within these cells (**Figure 8B**). The cell population was limited in number for those incubated with 16 mM NAC and H_2_O_2_ (**Figure 6**), so NAC could be having an intracellular cytoprotective effect on what little cells are present by reducing their intracellular ROS through elevating their total intracellular GSH content. These data helped to confirm the low end of the NAC cytoprotective window (*i.e.*, 0.25 mM and 1 mM NAC) and indicated the potential for high concentrations of NAC (*i.e.*, 4 mM and 16 mM NAC) to influence the total intracellular GSH level as a cytoprotective pathway against oxidative stress.

These data solely investigated the entire concentration of both GSH and GSSG present within the cells, however, the ability to measure both GSH and GSSG as individual concentrations through this assay is possible. This will be valuable future work to conduct as the intracellular GSH/GSSG ratio is also a measure of oxidative stress. Between the ROS and GSH data acquired already, NAC likely achieved its cell-mediated effects by reducing ROS and elevating total intracellular GSH levels at 4 mM and 16 mM NAC for H_2_O_2_-stressed NE-4Cs. As cells incubated with H_2_O_2_ and 0.25 - 16 mM GSH also expressed intracellular ROS, this is indicative of intracellular interactions between H_2_O_2_ and GSH. Most importantly, with the cell proliferation, cell viability, ROS, and GSH assessments for NE-4Cs, a therapeutic range for cytoprotection against oxidative stress was established as 2 - 8 mM NAC and 2 - 12 mM GSH.

### H_2_S, NAC, and GSH Neuroinductivity

In order to achieve significant healing in PNIs, therapeutic approaches must not only minimize neurotoxicity, but also facilitate neural stem cells to differentiate to regenerate tissue. To determine the ability SSMs have to induce neurogenesis, β3-tubulin was assessed as it is a primary neuronal marker that has been used to indicate neuronal differentiation [53]. A concentration four-fold below the cytotoxic limit for each molecule was chosen - 125 μM NaHS (H_2_S donor, 0.125 mM H_2_S), 4 mM NAC, and 4 mM GSH - to study SSM neuroinductivity. To evaluate their differentiative capacities, H_2_S, NAC, and GSH were evaluated against a negative control (*i.e.*, α-MEM media with no stimulus) for β3-tubulin expression over 7 days (**Figure 10**). The top panel for each image set is solely DAPI expression (*ex*: 359 nm, *em*: 457 nm), indicating the cell nuclei. The bottom panel for each image set is an overlay image of DAPI expression and Alexa Fluor® 647-antibody labeled β3-tubulin expression (*ex*: 488 nm, *em*: 507 nm). Each image represents triplicate samples for its treatment group. As immunofluorescent staining, on its own, is solely a qualitative method, corrected total cell fluorescence (CTCF) was also determined to provide semi-quantitative analysis of the data (**Figure 11**).

**Figure 10.**
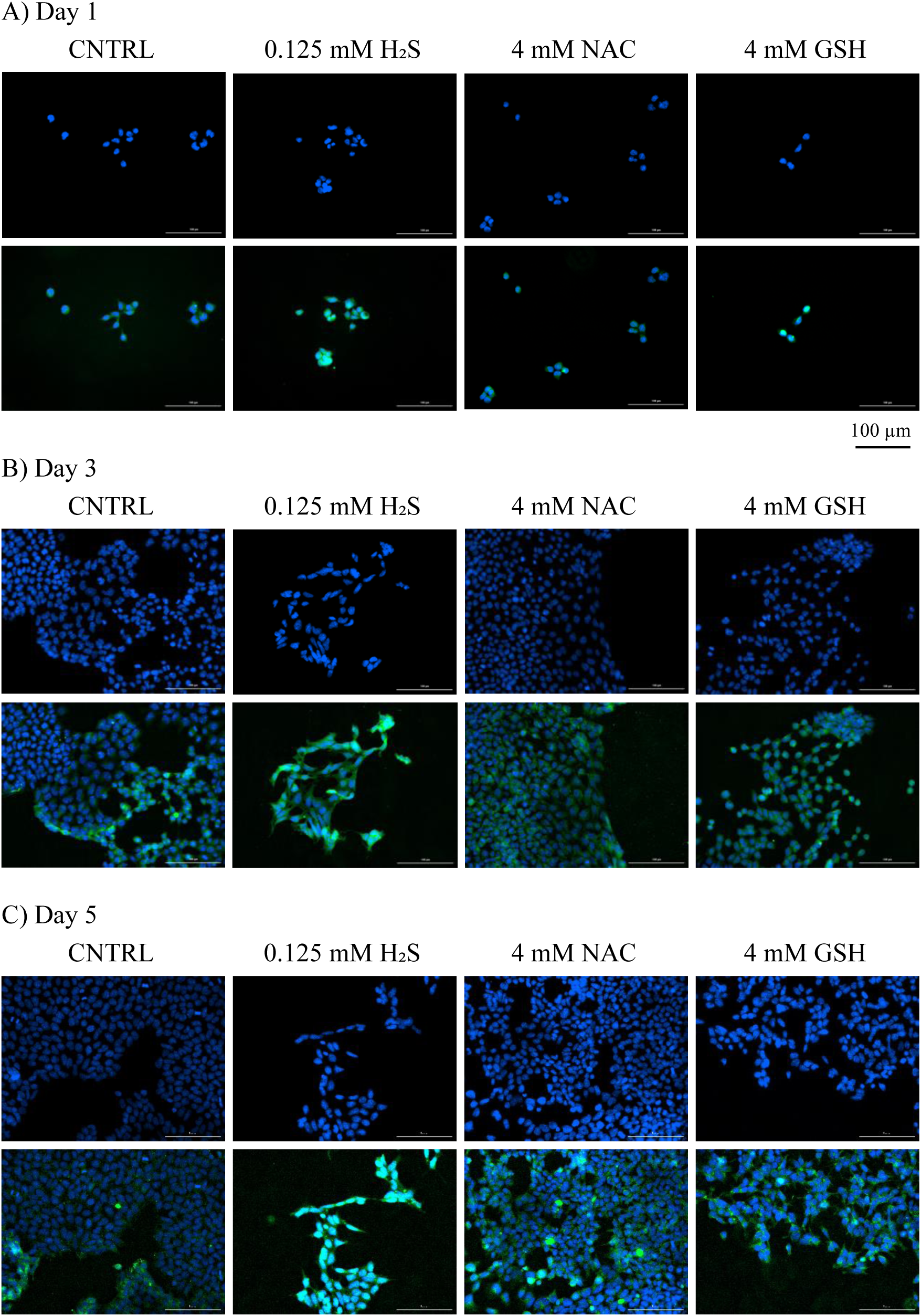
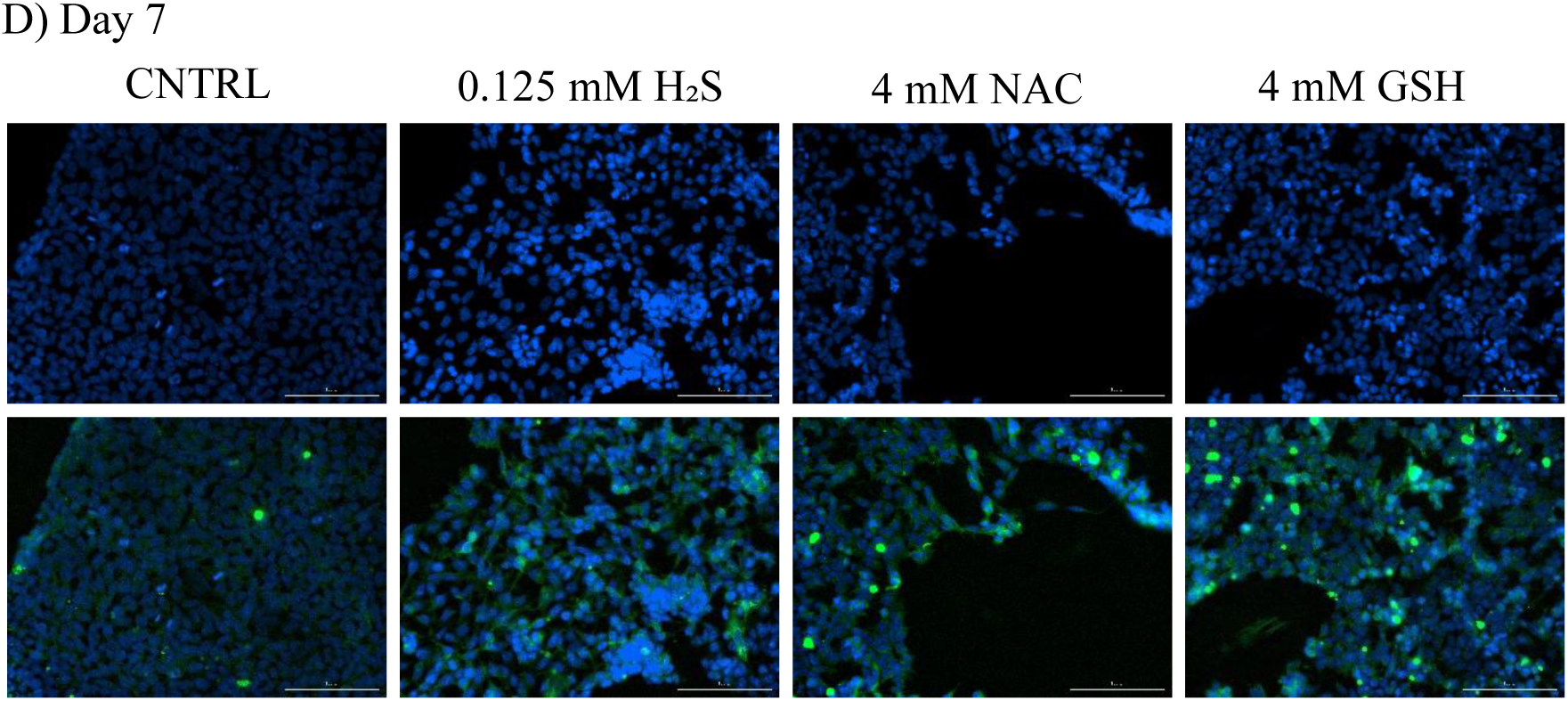
H_2_S, NAC, and GSH neuroinductivity in NE-4Cs. The ability for cells treated with α-MEM media with no stimulus, 0.125 mM H_2_S, 4 mM NAC, or 4 mM GSH to induce neural differentiation was assessed using nuclei and β3-tubulin expression staining. DAPI expression images are displayed on the top whereas DAPI + β3-tubulin expression overlay images are shown on the bottom at Day 1 (**A**), Day 3 (**B**), Day 5 (**C**) and Day 7 (**D**).

**Figure 11.**
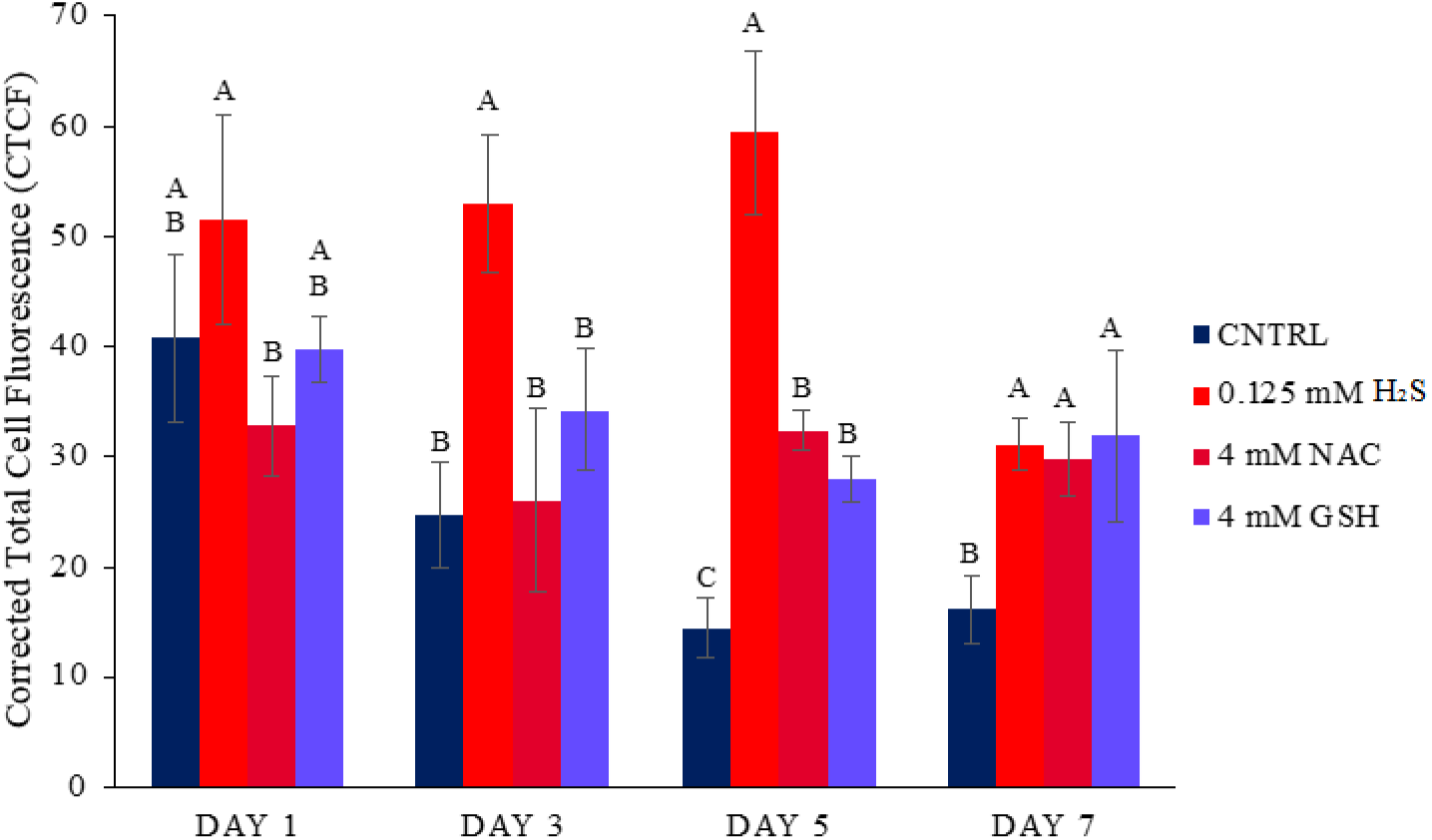
Corrected Total Cell Fluorescence (CTCF) of H_2_S, NAC, and GSH neuroinductivity in NE-4Cs. CTCF analysis was carried out for β3-tubulin expression for cells exposed to α-MEM culture media alone or with 0.125 mM H_2_S, 4 mM NAC, or 4 mM GSH as a semi-quantitative measure of NE-4C differentiation. Groups that possess different letters have statistically significant differences (p < 0.05) in mean whereas those that possess the same letter are statistically similar (N = 3).

Morphologically, elongated nuceli and neurite outgrowth are characteristics of neuronal differentiation [52]. At Day 1, the cells across all groups remained stem-like with no neurite outgrowths or elongated nuclei observed (**Figure 10A**). Although no morphological changes were identified, the CTCF data revealed some elevation in β3-tubulin for cells treated with H_2_S at Day 1, perhaps indicating some initial, but limited differentiation (**Figure 11**). By Day 3, morphological changes were detected in the H_2_S-treated cells when compared to the control cells and the cells subjected to NAC or GSH (**Figure 10B**). These cells possessed clearly elongated cell nuclei and neurite outgrowths between cells whereas all other groups had cells that persisted as circular and stem-like. This observation was supported by a statistically significant increase in CTCF for β3-tubulin expression for the cells treated with H_2_S (**Figure 11**). These observations were maintained into Day 5, with similar morphologies identified and CTCF values noted (**Figure 10C**). By Day 7, the morphological changes in the cells subjected to H_2_S could still be seen, but the β3-tubulin fluorescence visibly decreased (**Figure 10D**) which was also reflected in the fluorescence intensity data (**Figure 11**). As β3-tubulin is an early neuronal marker, it is possible that by Day 7, the decrease in fluorescence was due to a change in the differentiation state of the NE-4Cs. These findings demonstrated that H_2_S holds promise as a neuroinductive molecule whereas NAC and GSH did not exhibit the same influence on NE-4Cs. It should be noted that although some of the cells incubated with NAC and GSH had elevated β3-tubulin-related fluorescence intensity (**Figure 10**), the overall CTCF values were relatively similar to those of the control cells (**Figure 11**) and did not have some of the same morphological changes observed in the H_2_S-treated cells.

### Expanded H_2_S Neuroinductivity Window

To investigate the neuroinductivity of H_2_S further, a range of concentrations (*i.e.*, 7.75 - 500 μM H_2_S) were evaluated for their capacity to induce β3-tubulin expression in NE-4Cs. Fluorescent images of cells subjected to 7.75, 31, 125, and 500 μM H_2_S are provided in **Figure 12** as representative images of the whole data set of three images per treatment group. Similar to the previous imaging data, the top panel for each image set is only DAPI-labeled nuclei whereas the bottom panel has both DAPI staining as well as Alexa Fluor® 647-antibody labeled β3-tubulin expression after which CTCF was also calculated (**Figure 13**). None of the NE-4Cs showed any morphological changes at Day 1 (**Figure 12A**) which was supported by no statistically significant differences in CTCF data found across all cell treatments (**Figure 13**). However, by Day 3, there were neurite outgrowths and elongated nuclei within the H_2_S-treated cells as well as visible β3-tubulin expression throughout, regardless of H_2_S concentration (**Figure 12B**). These observations were also supported by the β3-tubulin CTCF data, as NE-4Cs subjected to 31 μM or 125 μM H_2_S exhibited significantly greater CTCF than the negative control cells while those incubated with 7.75 μM H_2_S and 500 μM H_2_S only had statistically significant, slightly elevated CTCF values (**Figure 13**). Similar findings were seen on Day 5, with continued morphological changes and β3-tubulin expression (**Figure 12C**). In fact, all H_2_S concentrations (*i.e.*, 7.75 - 500 μM H_2_S) conveyed significantly greater CTCF in treated cells than exposure to α-MEM media alone (**Figure 13**). By Day 7, β3-tubulin expression began to decrease for the higher H_2_S concentrations (*i.e.*, 31, 125, and 500 μM), however, there was still slight elevation in the CTCF of NE-4Cs incubated with H_2_S, especially in the lowest 7.75 μM H_2_S concentration (**Figure 12D** and **Figure 13**). This behavior could be indicative of lower H_2_S concentrations more gradually inducing differentiation after a prolonged exposure time; however additional studies would need to be conducted to investigate this phenomenon further.Based off these results, a neuroinductive window of 7.75 – 125 μM H_2_S was established as 500 μM H_2_S was found to be cytotoxic to NE-4Cs (**Figure 1A**). Cells treated within this range expressed β3-tubulin throughout the 7 days of the study, with greatest intensity at Days 3 - 7 dependent on the H_2_S concentration. It should be noted that these experiments did not titrate H_2_S low enough to show a loss of bioactivity, so it is possible this range could extend even lower, though the neuroinductive window established here is considerable and will able to be leveraged for future studies.

**Figure 12.**
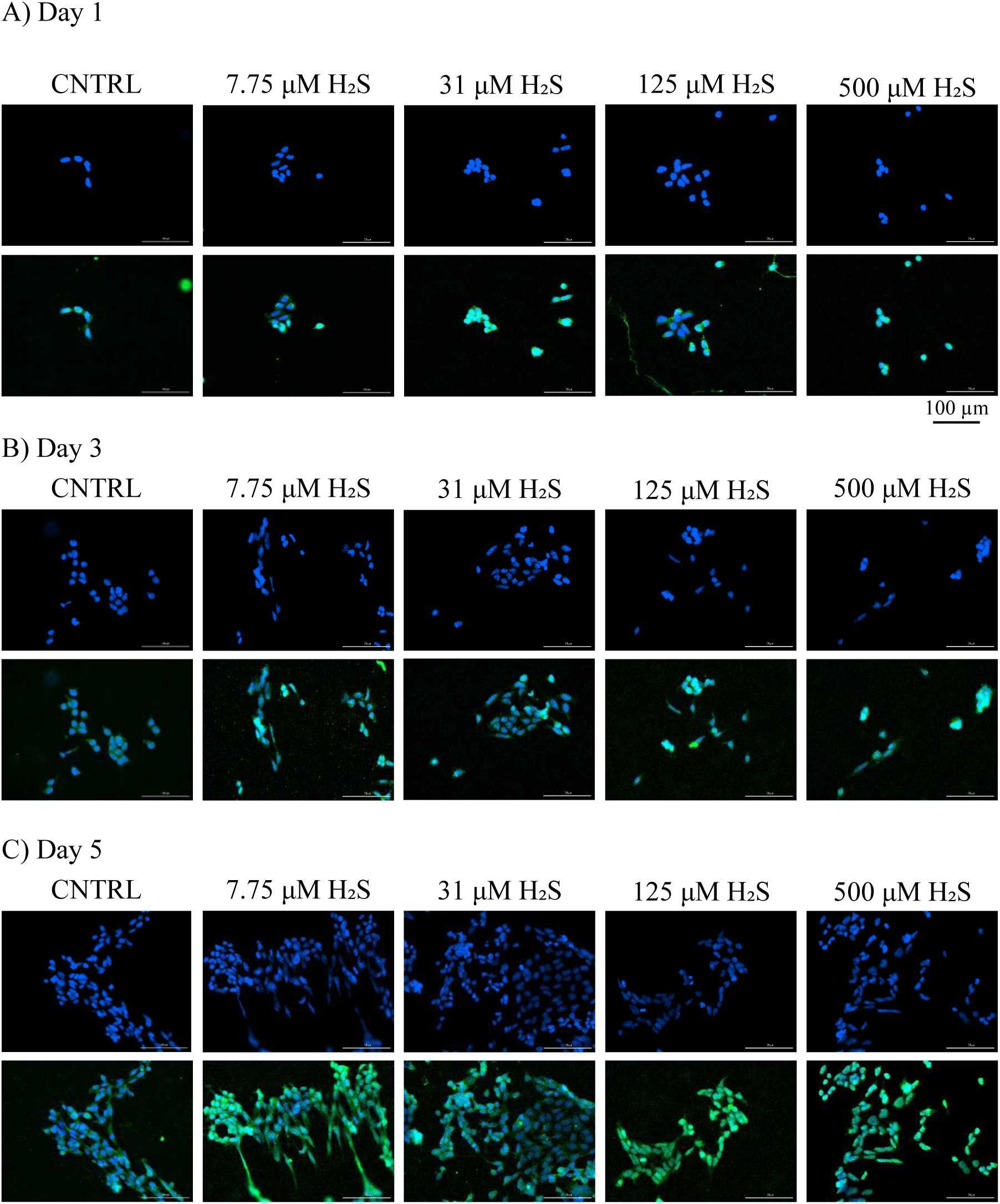
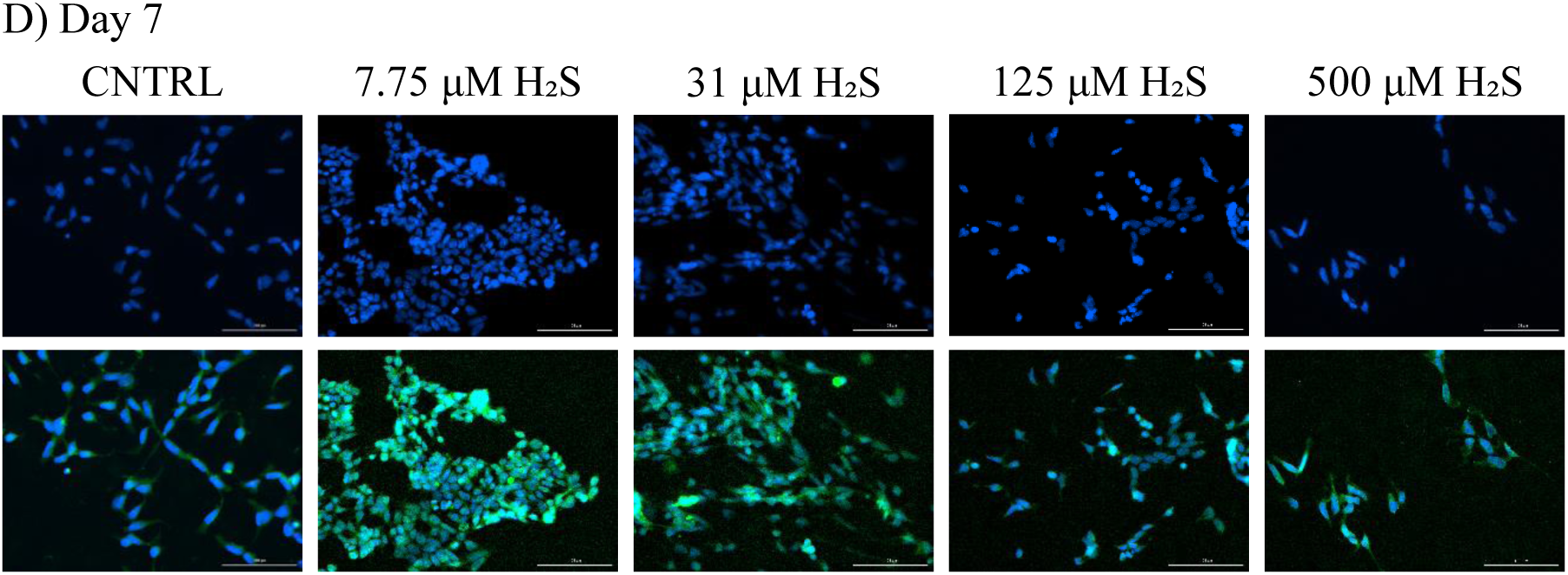
H_2_S neuroinductivity in NE-4Cs. The ability for cells treated with α-MEM media with no stimulus or 7.75 μM - 500 μM H_2_S to induce neural differentiation was evaluated using nuclei and β3-tubulin expression staining. DAPI expression images are displayed on the top whereas DAPI + β3-tubulin expression overlay images are shown on the bottom at Day 1 (**A**), Day 3 (**B**), Day 5 (**C**) and Day 7 (**D**).

**Figure 13.**
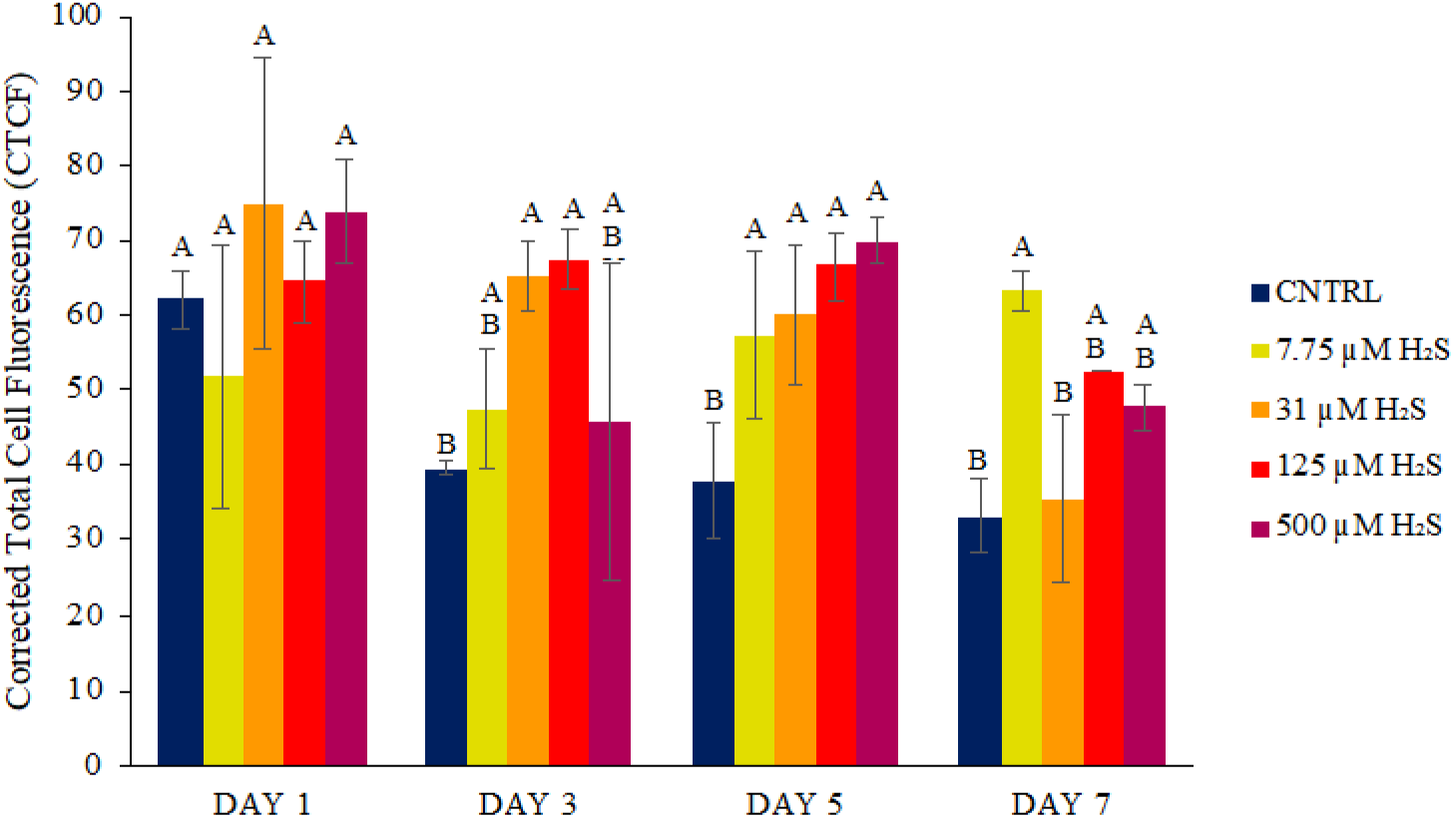
Corrected Total Cell Fluorescence (CTCF) of H_2_S neuroinductivity in NE-4Cs. CTCF analysis was conducted on the β3-tubulin expression for cells exposed to α-MEM culture media alone or with 7.75 μM - 500 μM H_2_S as a semi-quantitative measure of NE-4C differentiation. Groups that possess different letters have statistically significant differences (p < 0.05) in mean whereas those that possess the same letter are statistically similar (N = 3).

## CONCLUSIONS

This enclosed work was able to determine the respective therapeutics windows of NAC, GSH, and H_2_S. N-acetylcysteine (NAC) and glutathione (GSH) have been found to function as effective cytoprotectants mitigating reactive oxygen species (ROS) within cytoprotective windows spanning from 2 - 8 mM and 2 - 12 mM, for NAC and GSH, respectively. Hydrogen sulfide (H_2_S) exhibited neuroinductive properties in a now established neuroinductive window spanning from 7.75 - 125 μM, highlighting its role as a potential modulator of neuronal development and function. While promising, these SSMs, by themselves, are readily uptaken and/or easily diffuse away from one another in biological tissues thought their actions will likely require days to be effective for PNI repair. To address these concerns, controlled release of this suite of SSMs will need to be achieved. Many current treatment options, especially in oral or intravenous administration [53], employ ’burst’ release which results in peak concentrations followed by diminishing levels dropping below the therapeutic window [54]. The drawbacks of this delivery method include unpredictable toxicity risks due to rapid delivery [55] and the need for frequent dosing due to the short half-lives of the therapeutic paylods [54]. In contrast, controlled drug delivery strategies maintain more steady treatment levels, reducing administrations, ensuring consistent therapy, and minimizing overdose risks [54–57]. Specifically, delivering short-lived SSMs using degradable polymeric systems is advantageous for controlled drug delivery [56–58] and would enable the localized and sustained release of SSMs like H_2_S, NAC, and GSH, conveying a therapeutic response.

The simple molecular structures of H_2_S, NAC, and GSH lend to their direct incorporation into biodegradable polymers to achieve controlled release, an option often not available for larger molecules like proteins. As H_2_S is gaseous in nature, it requires a donor for effective delivery, one option of which is thioglutamic acid (GluSH, (**Figure 14A**). This analog of thioglycine and L- thiovaline reacts with bicarbonate to form N-carboxyanhydrides (NCAs) releasing H_2_S in the process [59–60]. By conjugating GluSH to another glutamic acid to form glutamylglutamic acid (GluGluSH), a diacid monomer can be generated that can be readily polymerized with itself to form a polyanhydride or with a diol to form a polyester, either of which can be used to facilitate prolonged, localized H_2_S release. A similar approach can be used to achieve controlled NAC release in that glutamic acid can be conjugated with NAC to form N-aceytlcysteinylglutamic acid (GluNAC, **Figure 14B**) which can also serve as a polymerizable monomer. Interestingly, GSH is already a diacid, so it requires no extra modification for polymer synthesis since it is readily able to be made into a polyanhydride (**Figure 14C**) or reacted with diols to generate polyesters. Future work with these novel SSM releasing monomers and polymers will include determining their therapeutic windows and modulating polymer chemistry to achieve cytoprotective and neuroinductive effects in neural stem cells for PNI repair applications in nerve guidance conduits.

**Figure 14.**
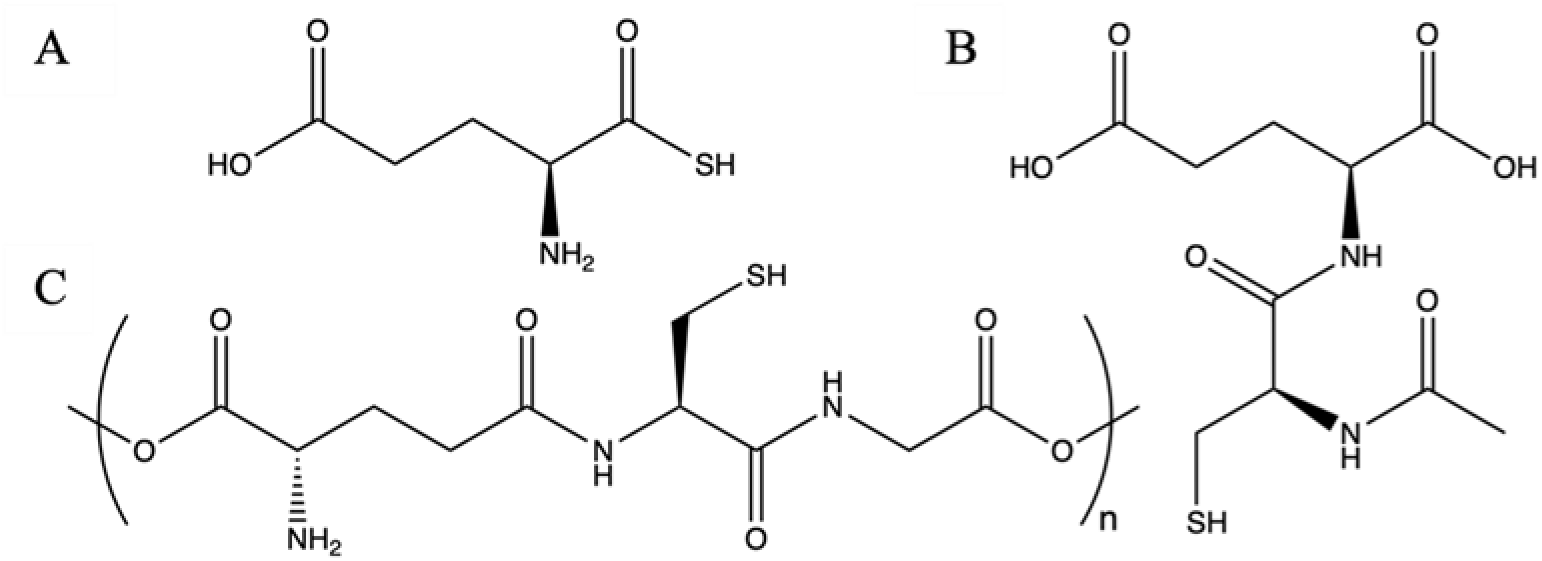
Thioglutamic acid (GluSH) (A), N-acetylcysteinylglutamic acid (GluNAC) (B) and Poly(glutathione anhydride) (Poly(GSHAn)) (C) chemical structures.

## STATEMENTS AND DECLARATIONS

### AUTHOR CONTRIBUTIONS

All authors participated in shaping the study’s conception and design. Kylie J. Dahlgren, August J. Hemmerla, Marissa A. Moore, and Daniela Calle were responsible for material preparation, data collection, and analysis. Kylie J. Dahlgren authored the initial draft of the manuscript, with subsequent drafts being written and edited by August J. Hemmerla and Bret D. Ulery. The final manuscript was reviewed and approved by all authors.

## REFERENCES

1. Taylor, C. A.; Braza, D.; Rice, J. B.; Dillingham, T. The Incidence of Peripheral Nerve Injury in Extremity Trauma. American Journal of Physical Medicine & Rehabilitation 2008, 87 (5), 381–385. 10.1097/phm.0b013e31815e6370.

2. Hakim, M.; N, K.; R, P.; D, T.; M, B.; H, H.; P, P.; A, F.; AD, W. A Review on Prevalence and Causes of Peripheral Neuropathy and Treatment of Different Etiologic Subgroups with Neurotropic B Vitamins. Journal of Clinical & Experimental Pharmacology 2019, 9 (4). 10.35248/2161-1459.19.9.262.

3. Grinsell, D.; Keating, C. P. Peripheral Nerve Reconstruction after Injury: A Review of Clinical and Experimental Therapies. BioMed Research International 2014, 2014, 1–13. 10.1155/2014/698256.

4. Wilhelm, J.; Xu, M.; Cucoranu, D.; Chmielewski, S.; Holmes, T.; Lau, K.; Bassell, G. J.; English, A. W. Cooperative Roles of BDNF Expression in Neurons and Schwann Cells Are Modulated by Exercise to Facilitate Nerve Regeneration. 2012, 32 (14), 5002–5009. 10.1523/jneurosci.1411-11.2012.

5. Rasappan, K.; Rajaratnam, V.; Wong, Y.-R. Conduit-Based Nerve Repairs Provide Greater Resistance to Tension Compared with Primary Repairs: A Biomechanical Analysis on Large Animal Samples. Plastic and Reconstructive Surgery. Global Open 2018, 6 (12), e1981. 10.1097/GOX.0000000000001981.

6. Schmidt, C. E.; Leach, J. B. Neural Tissue Engineering: Strategies for Repair and Regeneration. Annual Review of Biomedical Engineering 2003, 5 (1), 293–347. 10.1146/annurev.bioeng.5.011303.120731.

7. Kuihua, Z.; Chunyang, W.; Cunyi, F.; Xiumei, M. Aligned SF/P(LLA-CL)-Blended Nanofibers Encapsulating Nerve Growth Factor for Peripheral Nerve Regeneration. Journal of Biomedical Materials Research Part A 2013, 102 (8), 2680–2691. 10.1002/jbm.a.34922.

8. Richner, M.; Ulrichsen, M.; Elmegaard, S. L.; Dieu, R.; Pallesen, L. T.; Vaegter, C. B. Peripheral Nerve Injury Modulates Neurotrophin Signaling in the Peripheral and Central Nervous System. Molecular Neurobiology 2014, 50 (3), 945–970. 10.1007/s12035-014-8706-9.

9. Tajdaran, K.; Gordon, T.; Wood, M. D.; Shoichet, M. S.; Borschel, G. H. An Engineered Biocompatible Drug Delivery System Enhances Nerve Regeneration after Delayed Repair. Journal of Biomedical Materials Research Part A 2015, 104 (2), 367–376. 10.1002/jbm.a.35572.

10. Menorca, R. M.; Fussell, T. S.; Elfar, J. C. Nerve Physiology: Mechanisms of Injury and Recovery. Hand Clin. 2013, 29 (3), 317–330. DOI: 10.1016/j.hcl.2013.04.002.

11. Wang, J.-F.; Li, Y.; Song, J.-N.; Pang, H.-G. Role of Hydrogen Sulfide in Secondary Neuronal Injury. Neurochemistry International 2014, 64, 37–47. 10.1016/j.neuint.2013.11.002.

12. Massaad, C. A.; Klann, E. Reactive Oxygen Species in the Regulation of Synaptic Plasticity and Memory. Antioxid. Redox Signal. 2011, 14 (10), 2013–2054. DOI: 10.1089/ars.2010.3208.

13. Pizzino, G.; Irrera, N.; Cucinotta, M.; Pallio, G.; Mannino, F.; Arcoraci, V.; Squadrito, F.; Altavilla, D.; Bitto, A. Oxidative Stress: Harms and Benefits for Human Health. Oxid. Med. Cell. Longev. 2017, 2017, 8416763. DOI: 10.1155/2017/8416763.

14. Areti, A.; Yerra, V. G.; Naidu, V.; Kumar, A. Oxidative Stress and Nerve Damage: Role in Chemotherapy Induced Peripheral Neuropathy. Redox Biol. 2014, 2, 289–295. DOI: 10.1016/j.redox.2014.01.006.

15. Pun, P. B. L.; Lu, J.; Moochhala, S. Involvement of ROS in BBB Dysfunction. Free Radic. Res. 2009, 43 (4), 348–364. DOI: 10.1080/10715760902751902.

16. Jessen, K. R.; Mirsky, R. The Repair Schwann Cell and Its Function in Regenerating Nerves. J. Physiol. 2016, 594 (13), 3521–3531. DOI: 10.1113/JP270874.

17. Yowtak, J.; Lee, K. Y.; Kim, H. Y.; Wang, J.; Kim, H. K.; Chung, K.; Chung, J. M. Reactive Oxygen Species Contribute to Neuropathic Pain by Reducing Spinal GABA Release. Pain 2011, 152 (4), 844–852. DOI: 10.1016/j.pain.2010.12.034.

18. Solleiro-Villavicencio, H.; Rivas-Arancibia, S. Effect of Chronic Oxidative Stress on Neuroinflammatory Response Mediated by CD4+T Cells in Neurodegenerative Diseases. Front. Cell. Neurosci. 2018, 12, 114. DOI: 10.3389/fncel.2018.00114.

19. Jia, J.; Xiao, Y.; Wang, W.; Qing, L.; Xu, Y.; Song, H.; Zhen, X.; Ao, G.; Alkayed, N. J.; Cheng, J. Differential Mechanisms Underlying Neuroprotection of Hydrogen Sulfide Donors against Oxidative Stress. Neurochemistry International 2013, 62 (8), 1072–1078. 10.1016/j.neuint.2013.04.001.

20. Wang, M.; Tang, J.; Wang, L.; Yu, J.; Zhang, L.; Qiao, C. Hydrogen Sulfide Enhances Adult Neurogenesis in a Mouse Model of Parkinson’s Disease. Neural Regeneration Research 2021, 16 (7), 1353–1353. 10.4103/1673-5374.301026.

21. Gambari, L.; Grigolo, B.; Grassi, F. Hydrogen Sulfide in Bone Tissue Regeneration and Repair: State of the Art and New Perspectives. International Journal of Molecular Sciences 2019, 20 (20), 5231. 10.3390/ijms20205231.

22. Samuni, Y.; Goldstein, S.; Dean, O. M.; Berk, M. The Chemistry and Biological Activities of N-Acetylcysteine. Biochimica et Biophysica Acta (BBA) - General Subjects 2013, 1830 (8), 4117–4129. 10.1016/j.bbagen.2013.04.016.

23. Ezeriņa, D.; Takano, Y.; Hanaoka, K.; Urano, Y.; Dick, T. P. N-Acetyl Cysteine Functions as a Fast-Acting Antioxidant by Triggering Intracellular H_2_S and Sulfane Sulfur Production. Cell Chemical Biology 2018, 25 (4), 447–459.e4. 10.1016/j.chembiol.2018.01.011.

24. Liu, X.; Wang, L.; Cai, J.; Liu, K.; Liu, M.; Wang, H.; Zhang, H. N-Acetylcysteine Alleviates H2O2-Induced Damage via Regulating the Redox Status of Intracellular Antioxidants in H9c2 Cells. International Journal of Molecular Medicine 2019, 43 (1), 199–208. 10.3892/ijmm.2018.3962.

25. Pizzorno, J. Glutathione! *Integrative Medicine (Encinitas*, Calif.) 2014, 13 (1), 8–12.

26. Sedlak, T. W.; Paul, B. D.; Parker, G. M.; Hester, L. D.; Snowman, A. M.; Taniguchi, Y.; Kamiya, A.; Snyder, S. H.; Sawa, A. The Glutathione Cycle Shapes Synaptic Glutamate Activity. Proceedings of the National Academy of Sciences 2019, 116 (7), 2701–2706. 10.1073/pnas.1817885116.

27. Pandya, J. D.; Readnower, R. D.; Patel, S. N.; Yonutas, H. M.; Pauly, J. R.; Goldstein, G. D.; Rabchevsky, A. G.; Sullivan, P. F. N-Acetylcysteine Amide Confers Neuroprotection, Improves Bioenergetics and Behavioral Outcome Following TBI. 2014, 257, 106–113. 10.1016/j.expneurol.2014.04.020.

28. Won, S. J.; Kim, J.-E. .; Cittolin-Santos, G. F.; Swanson, R. A. Assessment at the Single-Cell Level Identifies Neuronal Glutathione Depletion as Both a Cause and Effect of Ischemia-Reperfusion Oxidative Stress. Journal of Neuroscience 2015, 35 (18), 7143–7152. 10.1523/jneurosci.4826-14.2015.

29. Muheremu, A.; Ao, Q. Past, Present, and Future of Nerve Conduits in the Treatment of Peripheral Nerve Injury. BioMed Research International 2015, 2015, 1–6. 10.1155/2015/237507.

30. Braga Silva, J.; Marchese, G. M.; Cauduro, C. G.; Debiasi, M. Nerve Conduits for Treating Peripheral Nerve Injuries: A Systematic Literature Review. Hand Surgery and Rehabilitation 2017, 36 (2), 71–85. 10.1016/j.hansur.2016.10.212.

31. Rahman, I.; Kode, A.; Biswas, S. K. Assay for Quantitative Determination of Glutathione and Glutathione Disulfide Levels Using Enzymatic Recycling Method. Nature Protocols 2006, 1 (6), 3159–3165. 10.1038/nprot.2006.378.

32. Zamaraeva, M. V.; Sabirov, R. Z.; Maeno, E.; Ando-Akatsuka, Y.; Bessonova, S. V.; Okada, Y. Cells Die with Increased Cytosolic ATP during Apoptosis: A Bioluminescence Study with Intracellular Luciferase. Cell Death & Differentiation 2005, 12 (11), 1390–1397. 10.1038/sj.cdd.4401661.

33. Pavel, M.; Renna, M.; Park, S. J.; Menzies, F. M.; Ricketts, T.; Füllgrabe, J.; Ashkenazi, A.; Frake, R. A.; Lombarte, A. C.; Bento, C. F.; Franze, K.; Rubinsztein, D. C. Contact Inhibition Controls Cell Survival and Proliferation via YAP/TAZ-Autophagy Axis. Nature Communications 2018, 9 (1), 1–18. 10.1038/s41467-018-05388-x.

34. Lu, M.; Hu, L.-F.; Hu, G.; Bian, J.-S. Hydrogen Sulfide Protects Astrocytes against H2O2-Inutamate Uptake. 2008, 45 (12), 1705–1713. 10.1016/j.freeradbiomed.2008.09.014.

35. Chadwick, W.; Zhou, Y.; Sung Soo Park; Wang, L.; Mitchell, N. A.; Stone, M. D.; Becker, K. G.; Martin, B.; Maudsley, S. Minimal Peroxide Exposure of Neuronal Cells Induces Multifaceted Adaptive Responses. PLOS ONE 2010, 5 (12), e14352–e14352. 10.1371/journal.pone.0014352.

36. Kimura, Y.; Kimura, H. Hydrogen Sulfide Protects Neurons from Oxidative Stress. The FASEB Journal 2004, 18 (10), 1165–1167. 10.1096/fj.04-1815fje.

37. Luo, Y.; Yang, X.; Zhao, S.; Wei, C.; Yin, Y.; Liu, T.; Jiang, S.; Xie, J.; Wan, X.; Mao, M.; Wu, J. Hydrogen Sulfide Prevents OGD/R-Induced Apoptosis via Improving Mitochondrial Dysfunction and Suppressing an ROS-Mediated Caspase-3 Pathway in Cortical Neurons. Neurochemistry International 2013, 63 (8), 826–831. 10.1016/j.neuint.2013.06.004.

38. Osborne, N. N.; Ji, D.; Shah, M.; Fawcett, R.; Sparatore, A.; Piero Del Soldato. ACS67, a Hydrogen Sulfide–Releasing Derivative of Latanoprost Acid, Attenuates Retinal Ischemia and Oxidative Stress to RGC-5 Cells in Culture. Investigative Ophthalmology & Visual Science 2010, 51 (1), 284–284. 10.1167/iovs.09-3999.

39. Lu, M.; Hu, L.-F.; Hu, G.; Bian, J.-S. Hydrogen Sulfide Protects Astrocytes against H2O2-Induced Neural Injury via Enhancing Glutamate Uptake. 2008, 45 (12), 1705–1713. 10.1016/j.freeradbiomed.2008.09.014.

40. Gao, C.; Chang, P.; Yang, L.; Wang, Y.; Zhu, S.; Shan, H.; Zhang, M.; Tao, L. Neuroprotective Effects of Hydrogen Sulfide on Sodium Azide-Induced Oxidative Stress in PC12 Cells. International Journal of Molecular Medicine 2017. 10.3892/ijmm.2017.3227.

41. Lu, M.; Hu, L.-F.; Hu, G.; Bian, J.-S. Hydrogen Sulfide Protects Astrocytes against H2O2-Induced Neural Injury via Enhancing Glutamate Uptake. 2008, 45 (12), 1705–1713. 10.1016/j.freeradbiomed.2008.09.014.

42. Zhou, D.; Shao, L.; Spitz, D. R. Reactive Oxygen Species in Normal and Tumor Stem Cells. Advances in cancer research 2014, 122, 1–67. 10.1016/B978-0-12-420117-0.00001-3.

43. Le Belle, J. E.; Orozco, N. M.; Paucar, A. A.; Saxe, J. P.; Mottahedeh, J.; Pyle, A. D.; Wu, H.; Kornblum, H. I. Proliferative Neural Stem Cells Have High Endogenous ROS Levels That Regulate Self-Renewal and Neurogenesis in a PI3K/Akt-Dependant Manner. Cell stem cell 2011, 8 (1), 59–71. 10.1016/j.stem.2010.11.028.

44. Ekshyyan, O.; Aw, T. Y. Decreased Susceptibility of Differentiated PC12 Cells to Oxidative Challenge: Relationship to Cellular Redox and Expression of Apoptotic Protease Activator Factor-1. Cell Death & Differentiation 2005, 12 (8), 1066–1077. 10.1038/sj.cdd.4401650.

45. Sung, Y.-J.; Cheng, C.; Chen, C.-S.; Huang, H.-B.; Huang, F.-L.; Wu, P.-C.; Shiao, M.- S.; Tsay, H.-J. Distinct Mechanisms Account for Beta-Amyloid Toxicity in PC12 and Differentiated PC12 Neuronal Cells. Journal of Biomedical Science 2003, 10 (4), 379–388. 10.1007/BF02256429.

46. Chen, L.; Liu, L.; Yin, J.; Luo, Y.; Huang, S. Hydrogen Peroxide-Induced Neuronal Apoptosis Is Associated with Inhibition of Protein Phosphatase 2A and 5, Leading to Activation of MAPK Pathway. The International Journal of Biochemistry & Cell Biology 2009, 41 (6), 1284–1295. 10.1016/j.biocel.2008.10.029.=

47. Aldini, G.; Altomare, A.; Baron, G.; Vistoli, G.; Carini, M.; Borsani, L.; Sergio, F. N-Acetylcysteine as an Antioxidant and Disulphide Breaking Agent: The Reasons Why. Free radical research 2018, 52 (7), 751–762. 10.1080/10715762.2018.1468564.48.

48. Bindoli, A.; Fukuto, J. M.; Forman, H. J. Thiol Chemistry in Peroxidase Catalysis and Redox Signaling. Antioxidants & Redox Signaling 2008, 10 (9), 1549–1564. 10.1089/ars.2008.2063.

49. Zamkova, M.; Khromova, N.; Kopnin, B. P.; Kopnin, P. Ras-Induced ROS Upregulation Affecting Cell Proliferation Is Connected with Cell Type-Specific Alterations of HSF1/SESN3/P21Cip1/WAF1pathways. Cell Cycle 2013, 12 (5), 826–836. 10.4161/cc.23723.

50. Wu, W.; Liu, B.; Xie, C.; Xia, X.; Zhang, Y. Neuroprotective Effects of N-Acetyl Cysteine on Primary Hippocampus Neurons against Hydrogen Peroxide-Induced Injury Are Mediated via Inhibition of Mitogen-Activated Protein Kinases Signal Transduction and Antioxidative Action. Molecular Medicine Reports 2018. 10.3892/mmr.2018.8699.

51. Aletta, J. M. Phosphorylation of Type III ?-Tubulin in PC 12 Cell Neurites during NGF-Induced Process Outgrowth. Journal of Neurobiology 1996, 31 (4), 461–475. 10.1002/(sici)1097-4695(199612)31:4%3C461::aid-neu6%3E3.0.co;2-7.

52. Ankam, S.; Teo, B. K. K.; Pohan, G.; Ho, S. W. L.; Lim, C. K.; Yim, E. K. F. Temporal Changes in Nucleus Morphology, Lamin A/c and Histone Methylation during Nanotopography-Induced Neuronal Differentiation of Stem Cells. Frontiers in Bioengineering and Biotechnology 2018, 6. 10.3389/fbioe.2018.00069.

53. Kim, S.; Kim, J.-H.; Jeon, O.; Kwon, I. C.; Park, K. Engineered Polymers for Advanced Drug Delivery. European Journal of Pharmaceutics and Biopharmaceutics 2009, 71 (3), 420–430. 10.1016/j.ejpb.2008.09.021.

54. Huang, X.; Brazel, C. S. On the Importance and Mechanisms of Burst Release in Matrix-Controlled Drug Delivery Systems. Journal of Controlled Release 2001, 73 (2-3), 121–136. 10.1016/s0168-3659(01)00248-6.

55. Bhowmik, D.; Gopinath, H.; Pragati Kumar, B.; Duraivel, S.; Sampath Kumar, K. THE PHARMA INNOVATION Controlled Release Drug Delivery Systems. 2012, 1.

56. Kikuchi, A.; Okano, T. Pulsatile Drug Release Control Using Hydrogels. Advanced Drug Delivery Reviews 2002, 54 (1), 53–77. 10.1016/s0169-409x(01)00243-5.

57. Langert, K. A.; Brey, E. M. Strategies for Targeted Delivery to the Peripheral Nerve. Frontiers in Neuroscience 2018, *12*. 10.3389/fnins.2018.00887.

58. Peltonen, S.; Alanne, M.; Peltonen, J. Barriers of the Peripheral Nerve. Tissue Barriers 2013, 1 (3), e24956. 10.4161/tisb.24956.

59. Zhou, Z.; von Wantoch Rekowski, M.; Coletta, C.; Szabo, C.; Bucci, M.; Cirino, G.; Topouzis, S.; Papapetropoulos, A.; Giannis, A. Thioglycine and L-Thiovaline: Biologically Active H_2_S-Donors. Bioorganic & Medicinal Chemistry 2012, 20 (8), 2675–2678. 10.1016/j.bmc.2012.02.028.

60. Kaur, K.; Enders, P.; Zhu, Y.; Bratton, A. F.; Powell, C. R.; Kashfi, K.; Matson, J. B. Amino Acid-Based H_2_S Donors: N-Thiocarboxyanhydrides That Release H_2_S with Innocuous Byproducts. Chemical Communications 2021, 57 (45), 5522–5525. 10.1039/D1CC01309B.

